# Intramacrophage RIL-seq uncovers an RNA antagonist of the *Salmonella* virulence-associated small RNA PinT

**DOI:** 10.1101/2025.04.07.647523

**Authors:** Hoda Kooshapour, Gianluca Matera, Elisa Venturini, Thorsten Bischler, Jörg Vogel, Alexander J. Westermann

## Abstract

*Salmonella* virulence chiefly relies upon two major pathogenicity islands, SPI-1 and SPI-2, which enable host cell invasion and intracellular survival, respectively. There has been increasing evidence for post-transcriptional control of SPI gene expression by Hfq-dependent small RNAs (sRNAs) such as PinT. This 80-nucleotide sRNA is highly expressed after *Salmonella* enters host cells and modulates the transition from the SPI-1 to SPI-2 program by targeting different virulence factor mRNAs. It has been elusive, however, how PinT activity could be counteracted when virulence gene suppression were to be relieved. To identify putative inhibitors of PinT, we have mapped the RNA interactome of *Salmonella* recovered from infected macrophages, using an optimized version of the RIL-seq method. Next to offering an unprecedented view of Hfq-mediated RNA interactions during *Salmonella*’s intracellular infection stage, RIL-seq uncovered the 3’ end-derived sRNA InvS as a direct negative regulator of PinT. Biochemical and genetic experiments suggest a decoy mechanism whereby InvS lifts the PinT-mediated repression of virulence factors. Additionally, InvS acts as an mRNA repressor of the host cell adhesion protein, MipA, and PinT interaction with InvS relieves *mipA* repression. Together, our work identifies a unique pair of antagonistic sRNAs in a growing post-transcriptional network of virulence gene regulation.

## INTRODUCTION

*Salmonella enterica* represents an important species of zoonotic and foodborne pathogens, responsible for a range of diseases in livestock and humans, from self-limiting gastroenteritis to potentially life-threatening typhoid fever in immunocompromised patients (1). Being a facultative intracellular pathogen, *Salmonella* can establish a replication niche inside epithelial and phagocytic host cells in the gut. *Salmonella*’s ability to infect mammals is mediated by intricate genetic systems, including the well-characterized *Salmonella* Pathogenicity Islands (SPIs) 1 and 2. These islands each encode an arsenal of secreted virulence effector proteins and the necessary type-III secretion systems (T3SS-1 and -2) to translocate these effectors into eukaryotic host cells (2,3). *Salmonella* infection is driven by sequential waves of expression of SPI-1 and SPI-2 genes, which underlie bacterial invasion and intracellular replication inside eukaryotic host cells, respectively (4,5). Numerous transcriptional regulators have been identified, that in their sum ensure SPI gene expression—which is costly and needs coordination with other cellular pathways—is only activated under infection-conducive conditions. However, there has also been growing evidence for a crucial role of post-transcriptional control in SPI gene expression, following observations that genetic inactivation of the global RNA-binding proteins CsrA, Hfq, and ProQ resulted in attenuated virulence (6–8). Relevant to the present study, Hfq is a conserved ∼60 kDa RNA-binding protein that primarily acts to mediate short base-pairing interactions of associated small regulatory RNAs (sRNAs) with their target mRNAs (9–12). *Salmonella* species encode ∼100 Hfq-associated sRNAs with diverse functions in bacterial physiology (13–17), including a proven or predicted regulation of virulence factors via mRNA targeting (14,18–25).

The most extensively characterized virulence-related sRNA in *Salmonella* species to date is PinT. Originally described as candidate STnc440 by sequencing Hfq-associated transcripts in *Salmonella enterica* serovar Typhimurium (13), PinT is an 80 nt-long sRNA from a horizontally acquired *Salmonella*-specific locus that also encodes RtsA, a co-activator protein of invasion genes (18). Its expression is positively regulated by PhoP/Q, the SPI-2-activating two-component system that is essential for intracellular survival (26,27). Moreover, dual RNA-seq profiling of different cell types has shown PinT to be the top induced sRNA as *Salmonella* invades and replicates in epithelial and macrophage host cells (18). Through repressing multiple mRNAs of specific SPI-1 and SPI-2 effectors, PinT influences the host response by modulating the inflammatory signaling pathways triggered during infection (18). A comprehensive transposon insertion sequencing screen linked *pinT* disruption to significant fitness defects in pigs and cattle, highlighting its role in host colonization (28). Over the past years, the PinT regulon has continuously been expanded, to now include seven direct target mRNAs, with mostly virulence-related functions (18,20,21). In sum, the previous work on PinT has established it as an exemplary sRNA that controls the temporal expression of two major virulence programs, thus facilitating the pathogen’s transition from host cell invasion to ensuring intracellular replication (18,29). However, it has remained a crucial unanswered question how the activity of this sRNA could be counteracted to eventually relieve PinT-mediated target (especially SPI-2; (21)) repression during later infection stages.

Given that both PhoP activity and PinT transcript levels remain high throughout infection, we considered it unlikely that a PinT antagonist would repress the sRNA at the level of synthesis (i.e., transcription), but would rather target the sRNA directly. Of note, several Hfq-dependent sRNAs have been found to be counter-regulated by RNA decoys or sponges, which often sequester the short ‘seed’ region through which the sRNA would otherwise interact with its target mRNAs (30–33). Therefore, we reasoned that potential PinT antagonists should show in RNA interactome maps produced by the popular RIL-seq (RNA interaction by ligation and sequencing) technique (34). RIL-seq predicts RNA-RNA interaction partners within bacterial cells through ligation on, and co-purification with Hfq, followed by high-throughput sequencing of the resulting chimeric fragments (35). A first application of RIL-seq to *Salmonella* grown in a SPI-1-inducing condition discovered OppX, an RNA sponge that acts antagonistically on the Hfq-associated sRNA and porin repressor, MicF, to modulate the cell envelope of extracellular *Salmonella* (30). However, this single *in vitro* condition unlikely captured all relevant RNA interactors of PinT, especially not those that act during the intracellular phase of the infection cycle.

This work pioneers the application of RIL-seq to bacteria growing inside eukaryotic cells, with the goal to experimentally identify an RNA antagonist of PinT activity. Through improving the sensitivity of the RIL-seq protocol, we were able to report the Hfq-associated RNA interactome of intra-macrophage *Salmonella.* Next to confirming previously predicted target mRNAs of PinT, our RIL-seq results predicted another horizontally acquired Hfq-associated sRNA, InvS, as key interactor of PinT. Follow-up characterization showed that InvS acts to sequester PinT, thereby relieving the repression of PinT target mRNAs. Additionally, RIL-seq suggested the *mipA* (a.k.a. *ompV*) mRNA, encoding an outer membrane protein involved in bacterial adhesion, as the primary target of InvS. Collectively, this study reveals a complex interplay of two Hfq-associated sRNAs and their targets in timing *Salmonella* virulence gene expression and adds virulence to the emergent theme of sRNA regulation by RNA decoys and sponges (36,37).

## RESULTS

### Establishment of RIL-seq for intramacrophage Salmonella

In order to globally map RNA-RNA interactions in an intracellular pathogen, we established Hfq RIL-seq for intra-macrophage *Salmonella* (Fig. 1A). As a major challenge, previous RIL-seq experiments were typically performed on >4×10^10^ bacterial cells (30,34), a quantity far too large to be reasonably recovered from host cells under standard infection conditions. Therefore, we systematically determined the minimal number of bacteria necessary to immunoprecipitate RNA with FLAG-tagged Hfq from *Salmonella* cells (Suppl. Fig. S1A). This showed that 10^7^ bacteria are sufficient for ready detection of Hfq by western blot, while a faint Hfq band was visible even for 10^6^ bacteria. To determine the infection conditions necessary to reach these cell numbers, we incubated murine RAW264.7 macrophages (an established host model (38,39)) with opsonized *Salmonella* for bacterial engulfment to occur. CFU enumeration of the recovered intracellular bacteria at defined time points suggested the minimally required bacterial cell number to be reached at 20 hours post-infection (p.i.) at a multiplicity of infection (MOI) of 50 (Suppl. Fig. S1B).

**Figure 1.**
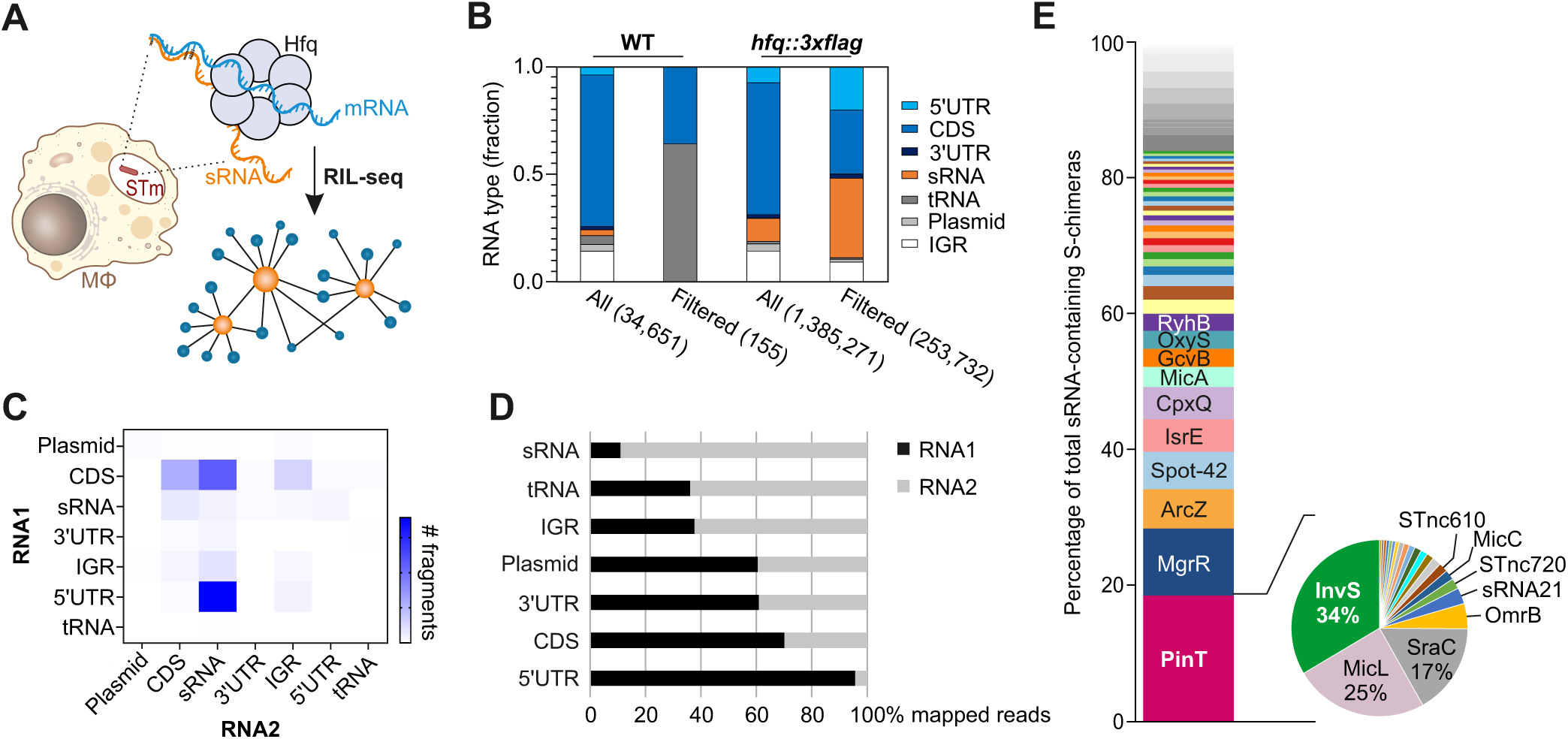
Intramacrophage RIL-seq workflow and global results. **A,** Cartoon representation of the intramacrophage RIL-seq procedure, from *Salmonella* inside RAW264.7 macrophages to the Hfq-associated RNA network depiction. **B,** Representation of the distribution of RNA types for significant chimeric fragments (number of fragments in S-chimeras; unfiltered [all] or filtered for ≥40 interactions) within the RIL-seq dataset derived from the wild-type and *hfq::3×FLAG* strains. UTR, untranslated region; CDS, coding sequence; sRNA, small RNA; tRNA, transfer RNA; plasmid, plasmid-encoded transcript; IGR, intergenic region. **C,** Two-dimensional heat map plotting the number of chimeric fragments corresponding to abundance-filtered S-chimeras according to the position of the RNA within the chimera (RNA1 = first read, RNA2 = second read). **D,** Distribution of chimeric fragments in RNA1 vs. RNA2 for each RNA type. **E,** The stacked bar chart denotes the relative abundance of sRNAs in S-chimeras. The pie chart to the right represents the total number of PinT interactions with various noncoding RNAs. The data plotted in panels B-E refer to unified values across both biological replicate intramacrophage RIL-seq experiments.

When subjecting the infection samples to RIL-seq, we found that direct lysis, without prior depletion of host-derived material (‘lysis strategy 1’ in Suppl. Fig. S1C), resulted in too few bacterial reads (∼10% of total reads; Supplementary Table 1). We therefore used a different harvesting protocol (40) in which the host cells are selectively lysed with a mild detergent, followed by selective centrifugation of the released bacteria (‘lysis strategy 2’ in Suppl. Fig. S1C). Although this extra step slightly reduced the amount of recovered Hfq protein (Suppl. Fig. S1D), it efficiently decreased the unwanted host read fraction and concomitantly increased the proportion of *Salmonella*-mapped reads to ∼34% (Supplementary Table 1). What is more, this bacterial enrichment prior to RIL-seq also yielded more chimeric reads (up from 1.7% with strategy 1 to almost 10% of the *Salmonella*-mapped reads with strategy 2; Supplementary Table 1). Based on these findings, we chose selective lysis (strategy 2) for our subsequent intramacrophage RIL-seq experiments.

### Intramacrophage RIL-seq predicts that PinT interacts with InvS sRNA

We performed intramacrophage Hfq RIL-seq in two biological replicates and sequenced the co-purified RNA to a depth of 60-130 million paired-end reads per library (Supplementary Table 1). Samples from macrophages infected with untagged Hfq (wild-type) bacteria served as controls. A global visualization of the datasets with regard to distribution and relative abundances of each RNA type was in accordance with previous Hfq RIL-seq studies of bacteria grown *in vitro* (30,35), such that sRNAs, mRNAs, and untranslated regions (UTRs) dominate the chimeras in the *hfq::3xFLAG* strain (Fig. 1B). As expected, most of the chimeric fragments were sRNA-coding sequence (CDS) and sRNA-mRNA 5’UTR chimeras (Fig. 1C). In addition, our *in*-*vivo* dataset recapitulated the previously described positional bias of CDSs and 5’UTRs to the first, and sRNAs to the second position of chimeras (30,34) (Fig. 1D). Reassuringly, the most abundant sRNAs on Hfq in intracellular *Salmonella* were PinT and MgrR (rank 1 and 2, with ∼18.5% or ∼9.8% of all sRNA-mapped reads, respectively), both of which are activated by the PhoPQ system (18,41) (Fig. 1E).

The intra-macrophage environment can be partially recapitulated *in vitro*, through batch growth of *Salmonella* in synthetic medium designed to mimic the vacuolar milieu, referred to as “SPI-2 medium” (42). For comparison with the above *in*-*vivo* data, we repeated our previous *Salmonella* RIL-seq study (30), yet in SPI-2 medium (Suppl. Fig. S2). A direct comparison between the sRNA distribution in the SPI-2 and the intramacrophage RIL-seq data showed an expected overall similarity (Suppl. Fig. S2D), with MgrR, PinT, and ChiX being the three top-enriched sRNAs, albeit their relative contribution to the chimeric fragments varied (compare Fig. 1E and Suppl. Fig. S2D). Whether these differences reflect the actual activities of specific sRNAs in the different conditions is unknown. While more chimeric fragments involving PinT were detected under SPI-2-inducing conditions compared to the intramacrophage environment, a substantial overlap was observed: 17 out of 25 targets detected under SPI-2 conditions were also found inside macrophages (Fig. S3), including several established PinT targets such as *grxA* (18) and *steC* (21). The overall agreement with respect to the most abundantly purified sRNAs and their interactomes validates the intramacrophage RIL-seq approach. Importantly, both datasets differed markedly from previous *Salmonella* Hfq RIL-seq data under a SPI-1-related condition (30) (Fig. S3C).

Given the dominance of PinT reads in both the intramacrophage and SPI-2 RIL-seq data, we reasoned that these datasets could point to potential negative regulators of this sRNA. Searching for PinT chimeras with other sRNAs, we predicted 19 (intramacrophage) or 13 (SPI-2) such putative PinT-sRNA pairs. The most frequent sRNA interactor of PinT in both conditions was InvS (pie chart in Fig. 1E, Suppl. Fig. S2D). Interestingly, InvS was also amongst the top-five transcripts in a previous MAPS study, in which we sequenced RNA interactors following co-purification with aptamer-tagged PinT sRNA from *Salmonella* grown in SPI-2 medium (21). Therefore, we chose to pursue InvS as a likely candidate of an RNA antagonist of PinT.

### InvS is a PhoP-induced 3’-derived sRNA and accumulates in intracellular Salmonella

InvS (a.k.a. STnc470) is a ∼90 nt-long, Hfq-associated sRNA (15,43). The InvS sequence is well conserved across *Salmonella* species (Suppl. Fig. S4A), but not present in other members of the Enterobacteriaceae. InvS has shown to promote *Salmonella* invasion of HeLa cells (44), yet its biogenesis and direct targets have remained elusive. InvS has been annotated as a 3’ end-derived sRNA since it overlaps with the 3’UTR of the coding gene *srfN* (a.k.a. *STM0082*; encoding a probable secreted protein). Given the absence of a predicted transcriptional start site (TSS) in the intergenic region between *srfN* and *invS*, yet presence of a predicted RNase E cleavage site coinciding with the adenine at the annotated 5’ end of the sRNA (45), we hypothesized that InvS is generated through ribonuclease processing of the polycistronic *SL0083A*–*srfN*–*invS* transcript (Fig. 2A). In support of this hypothesis, we observed that InvS stopped accumulating when the major endoribonuclease RNase E was inactivated in an established temperature-sensitive *Salmonella* RNase E mutant strain (*rne*^TS^; (46,47)) (Fig. 2B).

**Figure 2.**
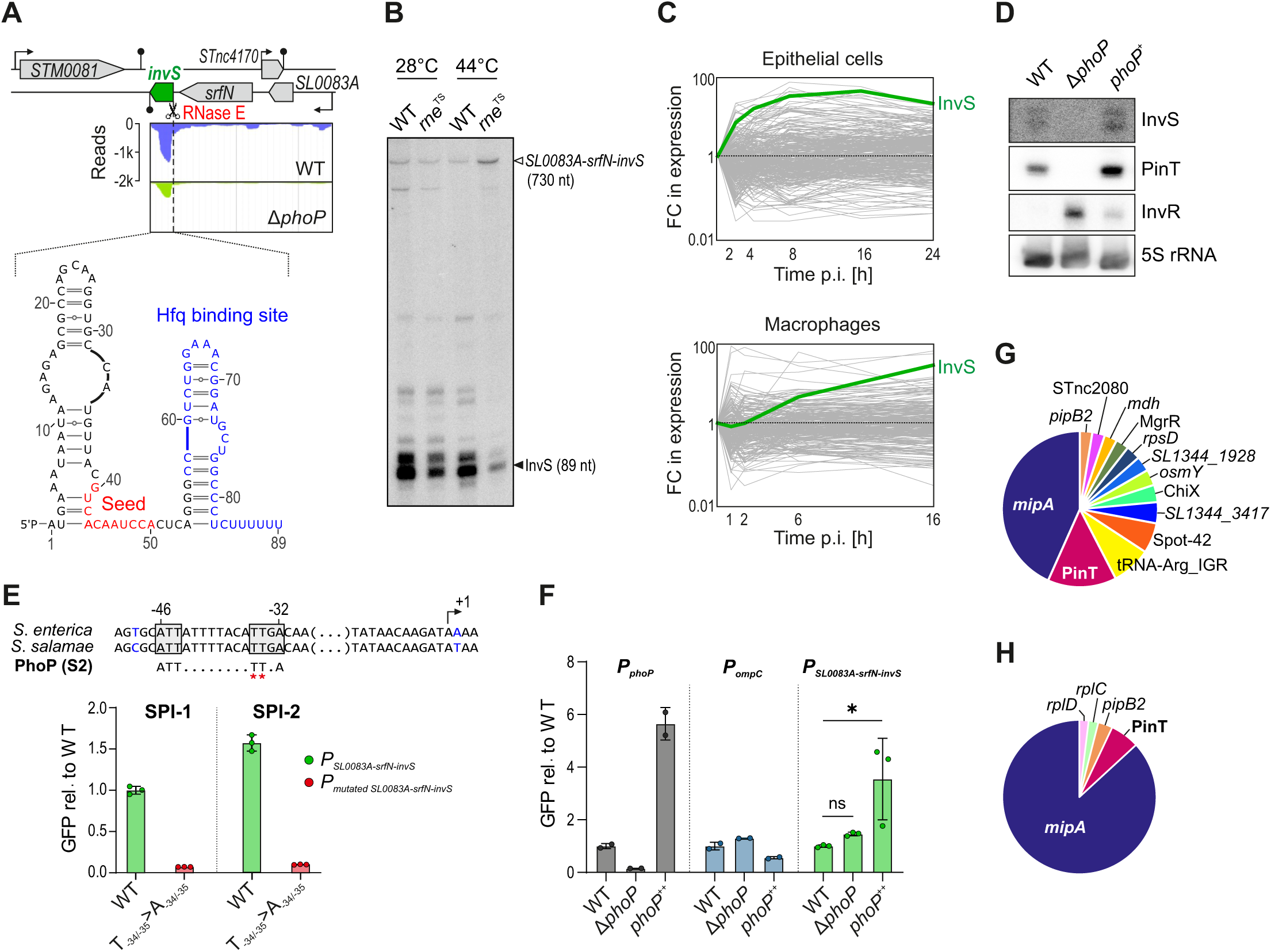
InvS expression, biogenesis, and interactome. **A,** InvS genomic location and structure. Expression of *SL0083A-srfN-invS* operon in a PhoP-deficient *Salmonella* mutant and its isogenic wild-type strain (48). **B,** InvS expression levels in a temperature-sensitive RNase E mutant (*rne*^TS^) and wild-type *Salmonella*. Northern blot showing RNA extracted from the respective overnight cultures grown at 28°C. Cultures were subsequently shifted to 44°C for 30 minutes before RNA purification. Both conditions are presented on the same blot for direct comparison. **C,** RNA-seq data from (18) shows InvS expression during infection of epithelial cells and macrophages relative to the corresponding expression levels in the bacterial inoculum. **D,** InvS expression follows that of the bona fide PhoP-induced sRNA PinT and is anti-correlated with SPI-1-associated InvR. **E,** PhoP box is required for transcriptional activation. A –138 to +162 fragment relative to the TSS of *SL0083A-srfN-invS* operon was cloned upstream of GFP on pAS093 plasmid, along with a mutant version containing point mutations (T-34/-35 → A-34/-35) on putative PhoP box. Constructs were transformed into WT and GFP expression in LB medium was measured by flow cytometry, showing the PhoP box is essential for activation of this operon. **F,** Validation of PhoP-dependent regulation using GFP reporter constructs. The GFP reporter construct from Fig. 2e was transformed into WT, Δ*phoP*, and *phoP*⁺⁺ strains (green). For comparison, GFP reporter constructs carrying the *phoP* promoter (positive control, black) and *ompD* promoter (negative control, blue) were also introduced into the same strains. The cultures grew in LB reaching OD=2.0 and the GFP levels were measured by flow cytometer. Significance was tested using one-way ANOVA, ns: *p* > 0.05, *: *p* ≤ 0.05. **G, H,** InvS interactome inside macrophages (G) and under SPI-2-inducing conditions (H), respectively, based on RIL-seq.

Interrogation of existing transcriptomics data (48) suggested reduced expression of the *SL0083A*–*srfN*–*invS* operon in a *phoP*-deficient *Salmonella* mutant strain relative to wild-type bacteria (Fig. 2A). In addition, while barely detectable in extracellular *Salmonella*, InvS strongly accumulated after invasion of epithelial cells or upon uptake of *Salmonella* by macrophages (18,49). In other words, InvS expression echoed that of the *bona fide* PhoP-induced sRNA PinT (Fig. 2C; Suppl. Fig. S4B). Using an *in*-*vitro* SPI-1/SPI-2 transition assay, we could reconstitute these expression profiles, again finding that InvS expression follows the kinetics of PinT induction (Suppl. Fig. S4C).

Northern blot probing confirmed that a Δ*phoP* mutation blunted InvS expression, while *trans*-complementation of the PhoP transcription factor (*phoP*^+^) fully restored InvS sRNA levels to that of the wild-type, both very similar to PinT (Fig. 2D). We also noticed a potential PhoP box (subclass S2 according to the classification of (50)) ∼35 bp upstream of the *SL0083A*–*srfN*–*invS* operon’s annotated TSS (Fig. 2E, upper). We thus generated a transcriptional reporter, cloning the region –138 to +162 relative to the TSS upstream of a GFP reporter cassette on a plasmid. A similar transcriptional fusion, yet with two point mutations in the putative PhoP box (T_-34/-35_ to A_-34/-35_), was made to serve as control. These reporters were inserted in wild-type *Salmonella* as well as in *phoP* deletion and overexpression strains. Using GFP fluorescent as a quantitative readout, these reporters revealed that the thymines at positions –34 and –35 are essential for transcriptional activation (Fig. 2E). Additionally, the data confirmed that PhoP enhances transcription of the *SL0083A*–*srfN*–*invS* operon; however, activation even occurred in the absence of PhoP, indicating additional regulatory factors are at play (Fig. 2F). Together, these data suggest that InvS biogenesis is dependent on both, PhoP-mediated transcriptional activation and RNase E-catalyzed terminal cleavage of the *SL0083A*–*srfN*–*invS* primary transcript.

### InvS represses the host-adhesion protein MipA by occluding its translation initiation region

RIL-seq predicted InvS as a major interactor of PinT, but would the reciprocal also be true? Intriguingly, when inspecting the InvS-harboring chimeras, PinT only scored second, with *mipA* mRNA being predicted as the most frequent RNA interactor in both the intramacrophage and SPI-2 datasets (Fig. 2G, H). The inferred interaction involved the region between nucleotides 40 and 50 of InvS (relative to its 5’ end; Fig. 2A) and the translational start codon of *mipA* (Fig. 3A, B). This would suggest that InvS acts as a translational repressor of the outer membrane protein, MipA. To test this hypothesis, we constructed a low-copy plasmid carrying the 5’ portion of *mipA* from the transcriptional start site to the 10^th^ codon (nucleotide positions −73 to +30 relative to the GUG start codon) fused to the open reading frame of superfolder green fluorescent protein (sfGFP, (51)) (Fig. 3C). Indeed, when introduced into *Salmonella*, this MipA-sfGFP translational fusion produced a high fluorescent signal, which was completely lost upon co-expression of the InvS sRNA from a second plasmid (Fig. 3D).

**Figure 3.**
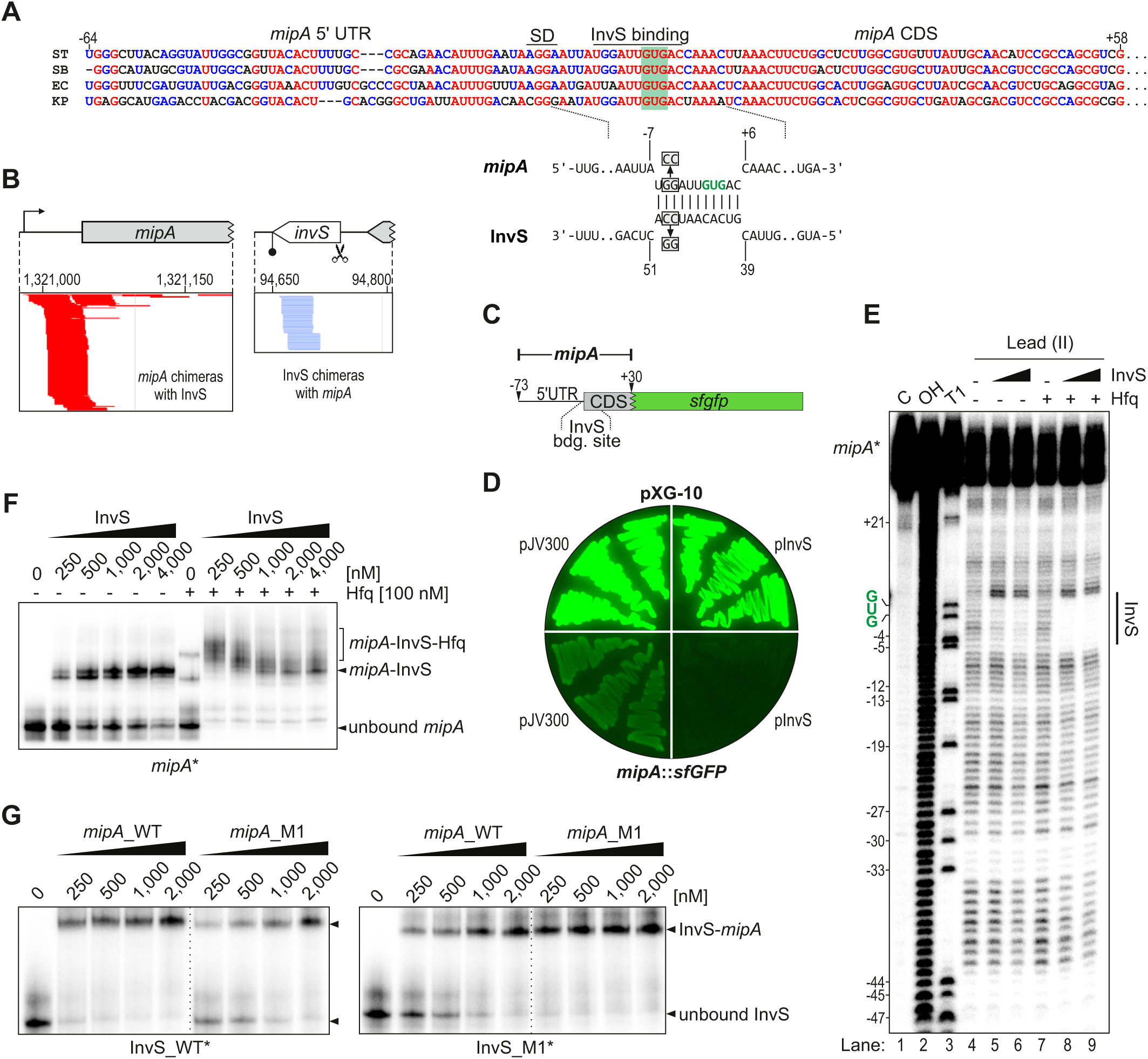
InvS represses MipA by occluding the translation initiation region of the mRNA. **A,** Upper: sequence alignment indicates that the InvS binding site on *mipA* mRNA is conserved among *S*. *enterica* servoar Typhimurium (ST), *S*. *bongori* (SB), *E*. *coli* (EC), and *K*. *pneumonia* (KP). SD refers to Shine-Dalgarno sequence. Numbers are relative to the translation start codon (+1 position). Lower: IntaRNA-predicted base-pair interaction between *Salmonella* InvS and the translation initiation region of *mipA*, with nucleotides mutated for panel G depicted in grey boxes. **B,** Genome browser-derived screenshot of the stacked reads of chimeric fragments mapping to the InvS sRNA and *mipA* mRNA (from replicate 1 of the intramacrophage RIL-seq). Reads corresponding to the first or second position in the chimera are colored red or blue, respectively. **C,** A low-copy plasmid was constructed to carry the *sfgfp* gene under the control of a region spanning the 5’UTR and the first 10 amino acids of *mipA* open reading frame (−73 to +30, relative to the start codon). This region was selected based on a predicted putative binding site for the InvS sRNA on the *mipA* mRNA. **D,** LB-agar plate showing the fluorescence of streaked strains containing the fluorescence control plasmid (pXG-10) with either overexpression of InvS (pInvS) or the empty vector (pJV-300) in the upper section and the corresponding strains containing the *mipA::sfgfp* fusion below. **E,** Structure probing experiment was performed using a 5′ end-radiolabeled *mipA* RNA fragment containing the predicted InvS-binding site. The fragment was incubated with increasing concentrations of unlabeled InvS sRNA, in the presence or absence of purified Hfq protein. **F,** EMSA with radioactively labelled *mipA* RNA fragment was performed to assess the binding interaction between InvS sRNA and the *mipA* transcript in absence and presence of Hfq. **G,** EMSA of radiolabeled InvS (wild-type [WT] or mutated [M1]) with increasing concentrations of the *mipA* mRNA fragment or a compensatorily mutated version thereof. The respective mutations are indicated in panel A, lower.

Next, we performed *in*-*vitro* biochemical assays to confirm the direct binding between InvS and the *mipA* translation initiation region. First, a structure probing experiment (Pb^2+^-induced cleavage) was conducted, incubating a 5’ end-radiolabeled *mipA* fragment containing the predicted InvS-binding site with increasing concentrations of unlabeled InvS in the presence or absence of purified Hfq protein. As predicted by RIL-seq, InvS clearly protected from cleavage at the region around the GUG start codon of *mipA* (Fig. 3E). While this protection was observed even without Hfq, the presence of this RNA chaperone in the reaction mix enhanced the intensity of the footprint. Second, Hfq also increased the binding affinity of InvS to the *mipA* mRNA fragment in an electrophoretic mobility shift assay (EMSA), with the dissociation constant (*k*_d_) decreasing from 1 µM without Hfq to <250 nM with Hfq (Fig. 3F). Importantly, the introduction of two point mutations (‘InvS_M1’; Fig. 3A) in the putative seed region of InvS reduced target binding, but binding was fully restored when introducing two compensatory mutations in the *mipA* fragment (Fig. 3G).

Collectively, these experiments suggest that InvS sequesters the start codon of the *mipA* mRNA and thereby inhibits MipA synthesis. While occurring even in the absence of an assisting protein under optimal conditions in a test tube, the InvS-*mipA* interaction is enhanced by Hfq.

### Seed region interaction of InvS and PinT reduces InvS stability

The second-ranked ligand of the InvS sRNA in the RIL-seq data was PinT (Fig. 2B, C). *In*-*silico* prediction using the IntaRNA online tool (52,53) suggested that InvS and PinT may form a duplex in which the putative seed region of either sRNA would be partially occluded (Fig. 4A). We therefore probed the predicted InvS-PinT interaction *in vitro*, incubating 5’ end-labeled PinT with increasing concentrations of unlabeled InvS (Fig. 4B). InvS protected PinT from Pb^2+^-induced cleavage, but only in presence of Hfq. Similarly, in an EMSA, 5’ end-labeled PinT was only shifted by InvS when the two RNAs were pre-incubated with Hfq (Fig. 4C). Collectively, these results imply that—at least under *in*-*vitro* conditions—the two Hfq-dependent sRNAs, InvS and PinT, form a complex on Hfq through recognizing each other’s seed sequence.

**Figure 4.**
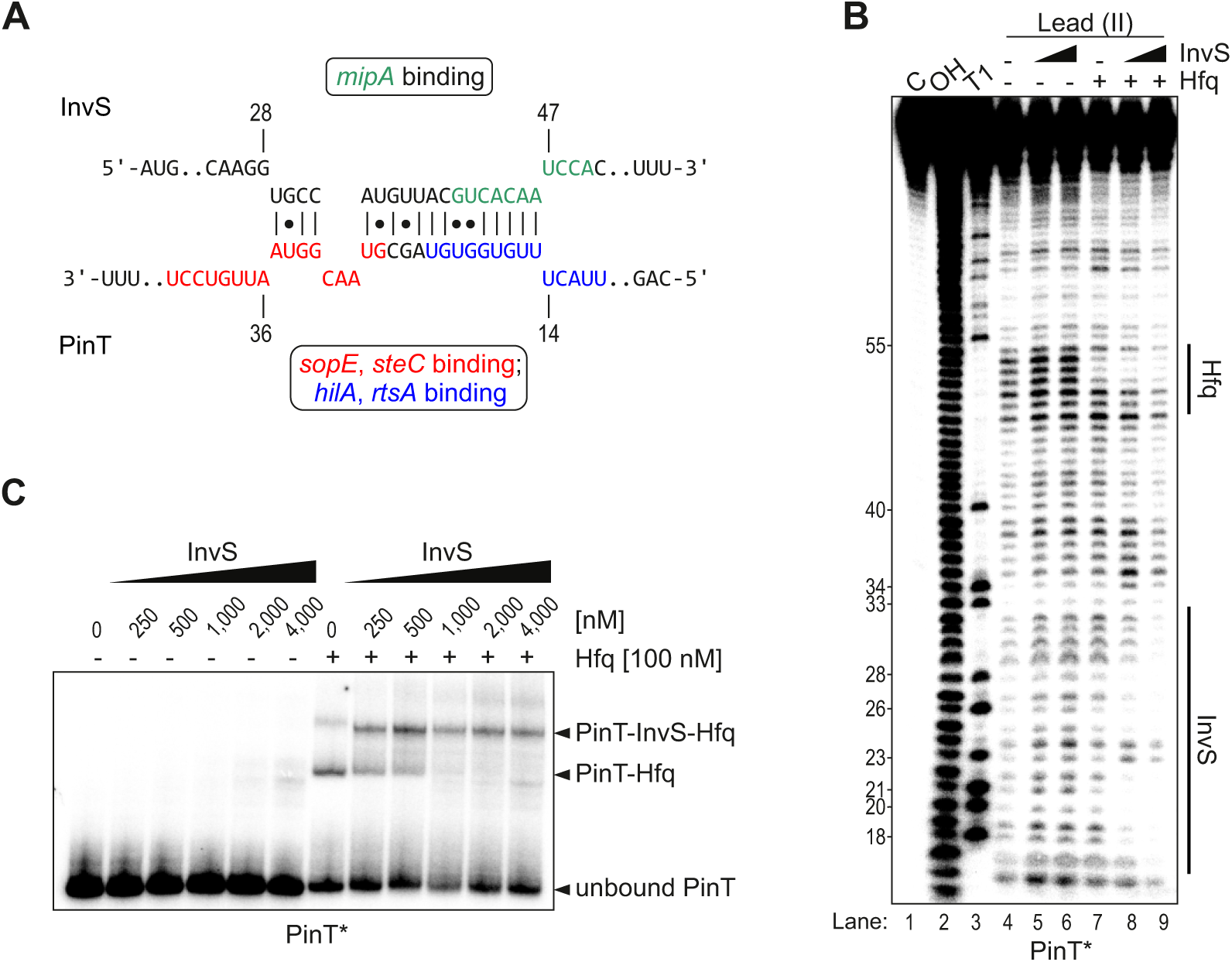
The two sRNAs InvS and PinT form an intermolecular duplex in the presence of Hfq. **A,** IntaRNA prediction of duplex formation between InvS and PinT. The respective seed regions of both sRNAs are colored. Note that PinT uses distinct seed regions to bind SPI-1 transcription factor mRNAs, *hilA* and *rtsA* (20) (blue), and to interact with effector-encoding mRNAs (18,21) (red), all of which are partially occluded within the InvS-PinT duplex. **B,** The protected PinT region corresponds to the predicted InvS binding site (see panel A). The second protected region near the 3’ end of PinT corresponds to the Hfq-binding site (15). **C,** Hfq facilitates the formation of a stable PinT-InvS complex. EMSA with 5′-end labeled PinT showed a stable complex with cold InvS, only after pre-incubation with Hfq, as indicated by a gel shift.

We next addressed the consequences of the PinT-InvS interaction *in vivo*, i.e., asking whether one sRNA regulated the other. For functional assays, we harnessed *Salmonella* strains lacking the *invS* or the *pinT* gene alone, or both (double knockout dubbed ‘ΔΔ’). In line with our previous findings (18), PinT did not influence *Salmonella* replication *in vitro* (Fig. 5A). Similarly, deletion of *invS* alone or in conjunction with *pinT* did not impact on *Salmonella* growth in any of these conditions. We also constructed strains overexpressing either PinT or InvS from a constitutive promoter on a high-copy plasmid. In LB medium, the PinT overexpression strain (*pinT*^++^) exhibited faster growth compared to the other strains, whereas InvS overexpression (strain *invS*^++^) slowed growth. In the SPI-2 medium, *pinT*^++^ had a longer lag-phase than the other strains, while both sRNA overexpression strains accumulated to higher cell densities than wild-type bacteria (Fig. 5A). Importantly, northern blot probing of total RNA extracted from the respective strains grown in SPI-2 medium showed elevated steady-state levels of InvS in the Δ*pinT* background, whereas PinT overexpression fully depleted InvS (Fig. 5B, left).

**Figure 5.**
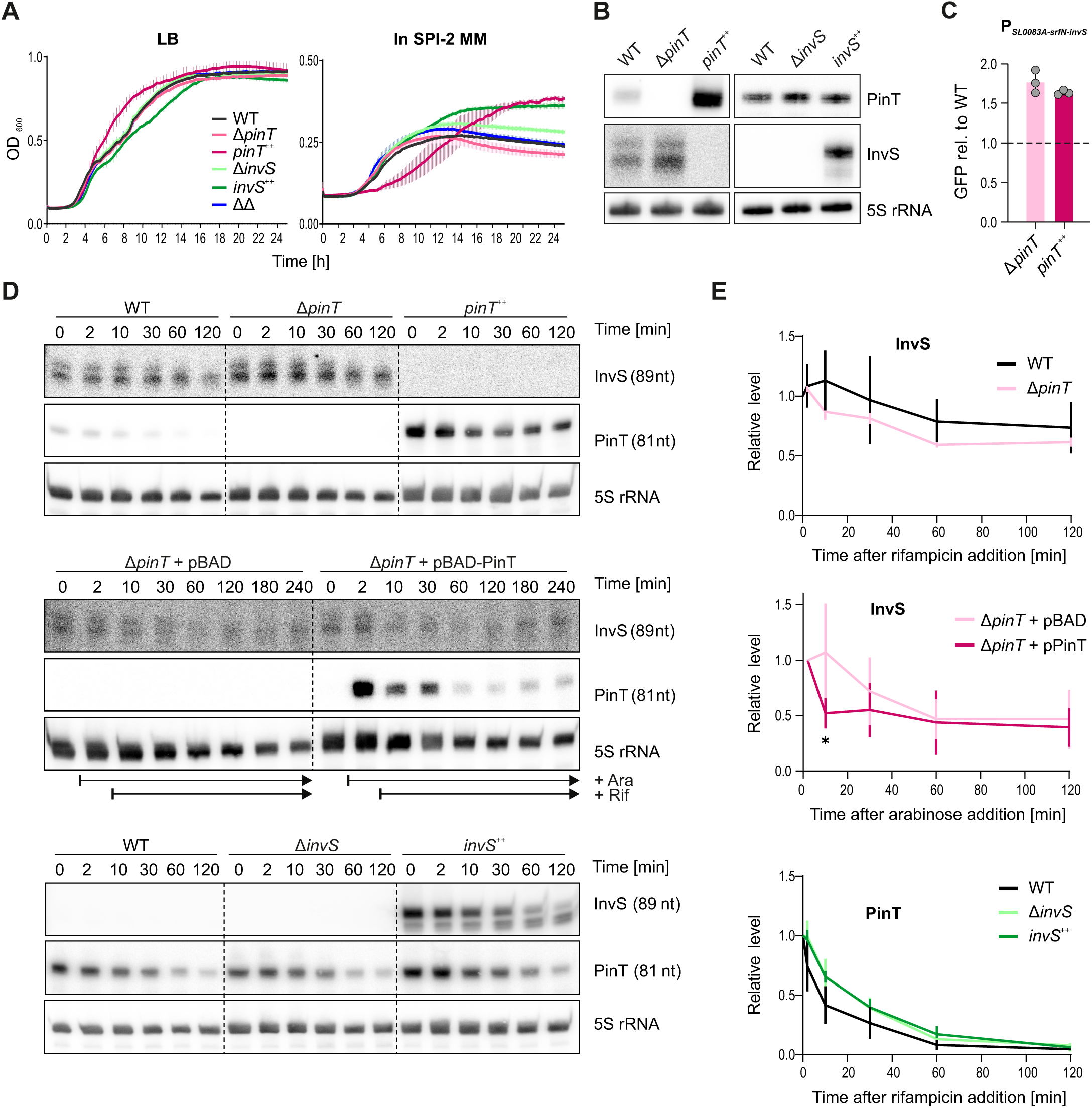
PinT reduces InvS stability. **A,** Growth curves of the indicated *Salmonella* sRNA deletion and overexpression strains relative to their parental wild-type under the SPI-1-(LB, OD_600_ 2.0) or SPI-2-inducing condition (SPI-2 minimal medium, OD_600_ 0.3). **B,** Northern blot-based measurement of InvS and PinT steady-state levels in the indicated strain backgrounds grown in SPI-2-inducing media. Three biological replicates per strain were grown in SPI-2 media and collected at OD_600_=0.3 **C,** Transcriptional reporter assay demonstrating that the transcriptional activity of the *SL0083A*-*srfN*-*invS* operon are independent of PinT levels. Data represent the mean ± SD from 3 biological replicates (individual gray dots). **D, E,** Rifampicin-mediated RNA-stability assay. At t_0_, *de*-*novo* transcription was halted and transcript decay traced over time and determined by northern blot. In each case, a representative northern blot out of 4 (upper blot), 3 (middle), or 6 (lower) independent replicate experiments is shown in panel D, and the corresponding quantifications in panel E. The asterisk indicates a statistically significant difference in relative InvS levels between the strain harboring the PinT expression plasmid and that containing the empty control vector at 8 min after halting *de*-*novo* transcription (*: *p* = 0.0268; unpaired t-test with Welch’s correction).

To explore the cause of PinT’s impact on InvS levels, we conducted several experiments. First, using transcriptional reporter strains, we ruled out a negative influence of PinT overexpression on the promoter that drives transcription of the *SL0083A*-*srfN*-*invS* locus (Fig. 5C). Second, rifampicin-mediated transcription runout experiments found no effect of deleting *pinT* on InvS stability (Fig. 5D, E, upper panels). As shown above, constitutive overexpression of PinT fully depleted InvS, this preventing half-life determination of the latter. We therefore opted for an inducible expression system, which allowed us to turn on PinT synthesis for two minutes prior to the addition of rifampicin. This revealed a PinT-dependent reduction of the half-life of InvS, as compared to an analogously treated strain harboring the empty control vector (Fig. 5D, E, middle panels). We concluded that PinT lowers InvS expression, at least partially through accelerated decay. In contrast, *invS* deletion or overexpression had little if any impact on PinT levels (Fig. 5B and lower panels in Fig. 5D, E), probably due to the relatively lower level of endogenous InvS compared to PinT. Obviously, the latter does not exclude the possibility that InvS may affect PinT activity; a hypothesis which we address below.

### PinT-suppressed virulence factors are de-repressed by InvS

We next evaluated potential effects of PinT and InvS on each other’s mRNA targets. Performing quantitative real-time PCR (qRT-PCR), we observed that all target mRNAs accumulated similarly in the different mutant strains and the wild-type in a SPI-1-inducing condition (LB, OD_600_ = 2.0), whereas expression changes were more pronounced in the SPI-2 medium, as expected given that both endogenous sRNAs are maximally induced in the latter medium (Fig. 6A). Particularly, absence of InvS decreased the levels of both tested PinT target mRNAs encoding the effector proteins SopE (18) and SteC (21). However, qRT-PCR failed to reliably quantify the levels of the InvS target, *mipA* mRNA. Therefore, we produced a polyclonal antiserum against endogenous MipA for immunoblotting. This antiserum, in combination with a previously constructed strain in which the C-termini of endogenous SopE (26.6 kDa) and SteC (44.3 kDa) are both FLAG-tagged in the chromosome (18), allowed us to detect all three proteins of interest in one sample. Western blot analysis under the SPI-1 and SPI-2 conditions revealed a slight increase in MipA protein levels by overexpressing PinT (Fig. 6B). InvS overexpression under the SPI-1 condition only entailed the downregulation of its cognate target protein, MipA, yet did not result in a de-repression of the PinT target proteins. In contrast, immunoblotting under the SPI-2-inducing condition (i.e. when endogenous PinT is highly expressed) revealed that overexpression of InvS increases the levels of both tested PinT targets at the protein level (Fig. 6B). In case of SteC, the positive effect of InvS was dependent on the presence of endogenous PinT, while for SopE, a slight increase was also visible in a Δ*pinT* background.

**Figure 6.**
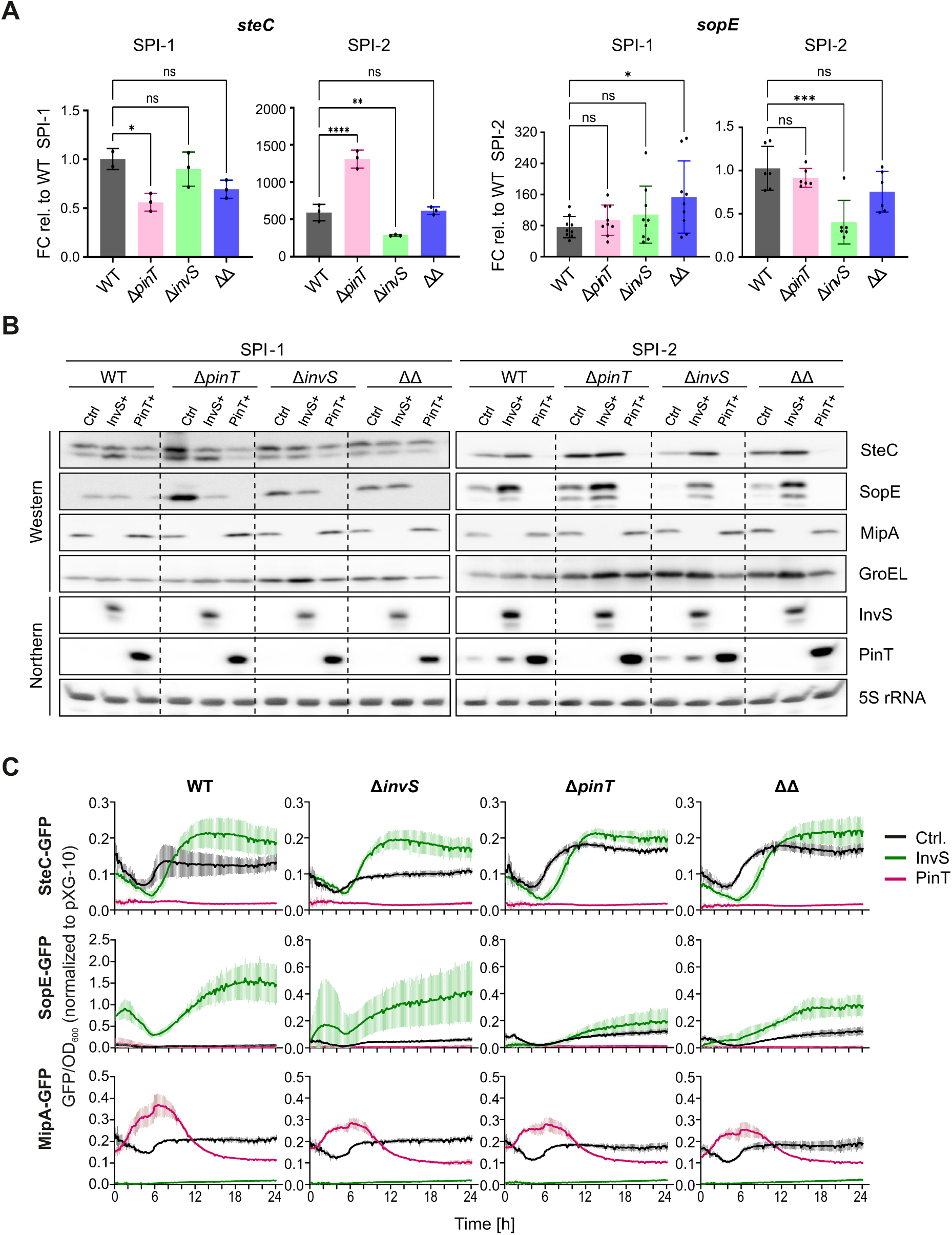
PinT and InvS mutually affect each other’s target genes. **A,** qRT-PCR analysis of PinT target levels. Quantitative real-time PCR (qRT-PCR) was performed to measure the expression levels of *sopE* and *steC* (SPI-1 and SPI-2 effector proteins respectively) in WT, Δ*pinT*, Δ*invS*, and ΔΔ strains. The RNA samples were collected under SPI-1 (LB, OD=2.0) and SPI-2 (SPI-2 MM, OD=0.3) inducing conditions. The quantification cycle (Cq) values were normalized to those of the reference gene 5S rRNA. Data represents the mean ± SD from ≥ 3 biological replicates (shown on bars by black dots), each with ≥ 3 technical replicates. Statistical significance was determined using one-way ANOVA, ns: *p* > 0.05, *: *p* ≤ 0.05, **: *p* ≤ 0.01, ***: *p* ≤ 0.001, ****: *p* <0.0001. **B,** Western blot analysis of endogenous MipA, SopE-FLAG, and SteC-FLAG protein levels. Western blot was performed using a custom antibody raised against endogenous MipA (24 kDa) and a strain with chromosomally FLAG-tagged C-termini of endogenous SopE (26.6 kDa) and SteC (44.3 kDa). Protein samples were collected from SPI-1 and SPI-2 inducing conditions and separated by SDS-PAGE, followed by immunoblotting with anti-MipA and anti-FLAG antibodies. Data shown are representative of three independent biological replicates. GroEL was used as a loading control and the integrity of the strains was tested by northern blots. **C,** Dual plasmid reporter assay of InvS and PinT targets in defined mutant backgrounds. Each background harbors an InvS or a PinT expression plasmid, or the empty vector control. Data derive from three biological and two technical replicates. OD_600_ and fluorescence intensities were read out in a plate reader for 24 hours. GFP fluorescence was first normalized by calculating the GFP/OD₆₀₀ ratio for each well (Suppl. Fig. S6), followed by second normalization to the mean GFP/OD ratio of the pXG10 control (set to 1).

Hence, we asked whether InvS could also directly target the *sopE* mRNA. Although no *sopE*-InvS chimeras were detected under any of the tested RIL-seq conditions (Fig. 2C, D) (30), a targeted *in*-*silico* analysis suggested potential base pairing of InvS with the translation initiation region of *sopE*, through the same region that recognizes *mipA* (Supplementary Fig. S5A). However, EMSAs of a radioactively labeled *sopE* 5’ fragment incubated with increasing concentrations of InvS did not support such interaction, regardless of the presence of Hfq (Supplementary Fig. S5B). We then assessed if InvS could titrate Hfq away from PinT-*sopE* RNA duplexes. Performing competitive EMSAs in the presence of Hfq, we indeed observed increasing InvS concentrations to sequester Hfq. This was accompanied by the depletion of the Hfq protein from its complex with PinT and *sopE* and an increase of the fractions of PinT-*sopE* dimers and free *sopE* mRNA (Supplementary Fig. S5C). We conclude that the positive influence of InvS on the targets of PinT is mostly through targeting PinT, although we cannot rule out the possibility that titration of Hfq away from the PinT-target mRNA duplexes may contribute to relieving target repression.

Lastly, to provide a temporal angle to the inferred regulations, we harnessed existing translational reporter constructs based on sfGFP. *Salmonella* strains harboring either MipA-sfGFP, SopE-sfGFP or SteC-sfGFP were grown for 24 hours in SPI-2 medium and cell density (OD_600_) and fluorescence intensity measured in 10 minutes intervals (Fig. 6C; Suppl. Fig. S6). The above MipA reporter (Fig. 3C) was again strongly repressed by overexpression of InvS (Fig. 6C). PinT overexpression accelerated the accumulation of MipA-sfGFP, but this effect was seemingly independent of endogenous InvS. SteC-sfGFP fluorescence was repressed by PinT (Fig. 6C), as expected (21). In line with the observations made on mRNA and protein levels in the SPI-2 medium (Fig. 6A, B), InvS induction led to the de-repression of SteC, with the effect being more pronounced in the presence of endogenous PinT (Fig. 6C). Lastly, SopE-sfGFP accumulated when InvS was overexpressed or *pinT* deleted (Fig. 6C). Again, the positive influence of InvS on SopE was markedly reduced in the absence of endogenous PinT.

Altogether, these observations imply that the interaction of InvS and PinT leads to a reduction in the levels of the lesser abundant InvS sRNA. Additionally, high levels of InvS result in the sequestration of PinT and consequently, elevate the protein levels of the direct PinT targets SopE and SteC. PinT overexpression, in turn, changes the kinetics of MipA production, accelerating the accumulation of this InvS target.

## DISCUSSION

Along our quest to characterize the function of *Salmonella* virulence-related sRNAs, we have been focusing on PinT (54). This sRNA is transcriptionally activated upon host cell entry through the PhoPQ two-component system (18) and represses translation of several regulatory and effector proteins of the SPI-1 and SPI-2 regulons (18,20,21). However, how PinT-mediated SPI-2 repression would be relieved at later stages of infection, when this virulence program is fully activated, has remained unknown. We previously speculated that intracellularly, PinT might eventually be sponged by (an)other Hfq-associated sRNA(s), thereby mitigating post-transcriptional SPI-2 inhibition (21). However, the identification of such a postulated PinT antagonist has been hindered by a shortage of suitable methods.

RIL-seq is a powerful tool that provides a comprehensive view on a bacterium’s post-transcriptional regulatory network. Since its establishment in *E*. *coli* (34), the method has been continuously improved and adapted to various bacterial species and experimental conditions (31,32,34,55–58). However, the relatively high input requirements have so far limited the application of the technique to large-scale *in*-*vitro* experiments, which fail to fully replicate the complexity of natural bacterial host environments. For example, facultative intracellular pathogens such as *Salmonella* Typhimurium enter different mammalian host cell types, wherein they create an intracellular replication niche. Despite the relevance of this intracellular *Salmonella* subpopulation for disease development (59), the global RNA-RNA interactome has not previously been charted during the intracellular stage of this—or any other—pathogen’s infection cycle. Here, by systematically scaling down bacterial cell numbers, we reduced the input material from 4×10^10^ bacteria in conventional RIL-seq to 10^6^. Thanks to this 10,000-fold increase in sensitivity, we managed to capture the Hfq-associated RNA interactome of intramacrophage *Salmonella* purified from infected macrophages. *Salmonella* represents a suitable model organism to establish intramacrophage RIL-seq as defined growth media exist (42,60) that can mimic certain—albeit not all (49)—cues of the milieu of its *in vivo* niche, to serve as a reference for intramacrophage RIL-seq. Reassuringly, we observed overall large similarity between our intramacrophage and SPI-2 datasets.

Interrogation of the intramacrophage RIL-seq data for PinT-interacting sRNAs highlighted InvS, an sRNA with an established virulence phenotype (44), whose molecular function was elusive. According to our RIL-seq data, the primary target of InvS was the mRNA encoding MipA (a.k.a. OmpV). Of note, MipA has an established role in mediating adhesion of *Salmonella* to epithelial cells (61). We demonstrated here that InvS represses MipA protein synthesis by annealing to the corresponding mRNA and interfering with translation initiation. This seemingly creates a conundrum: how come *Salmonella* Δ*invS* mutants (that produce more MipA) are less invasive than wild-type bacteria (44)? Previously, InvS was shown to activate the expression of several SPI-1 genes, yet none of the corresponding transcripts seemed to be directly targeted by this sRNA (44). Here, we found that the second ranked InvS interactor was the PinT sRNA, a known negative regulator of SPI-1 and SPI-2 genes (18,20). We showed that InvS expression relieves PinT-mediated repression of both a SPI-1 (*sopE*) and SPI-2 (*steC*) target transcript. The positive influence of InvS on those virulence factors was partially dependent on endogenous PinT. Based on these collective findings, we conclude that InvS antagonizes PinT-mediated virulence factor repression.

*Salmonella* sRNAs may act as molecular timers of proper virulence program activation along the pathogen’s infection cycle (18–20,22,23,62). For example, PinT counteracts SPI-1-mediated invasion and delays SPI-2 onset (18,20,21). As we have uncovered here, InvS also belongs to this group of sRNA timers, as it represses the MipA adhesion protein, while promoting invasion through the activation of T3SS-1 effector translocation (44), thereby likely facilitating the shift from host cell adhesion to invasion. Importantly, since InvS and PinT antagonize one another, the regulatory networks of these two molecular timers are interconnected (Fig. 7).

**Figure 7:**
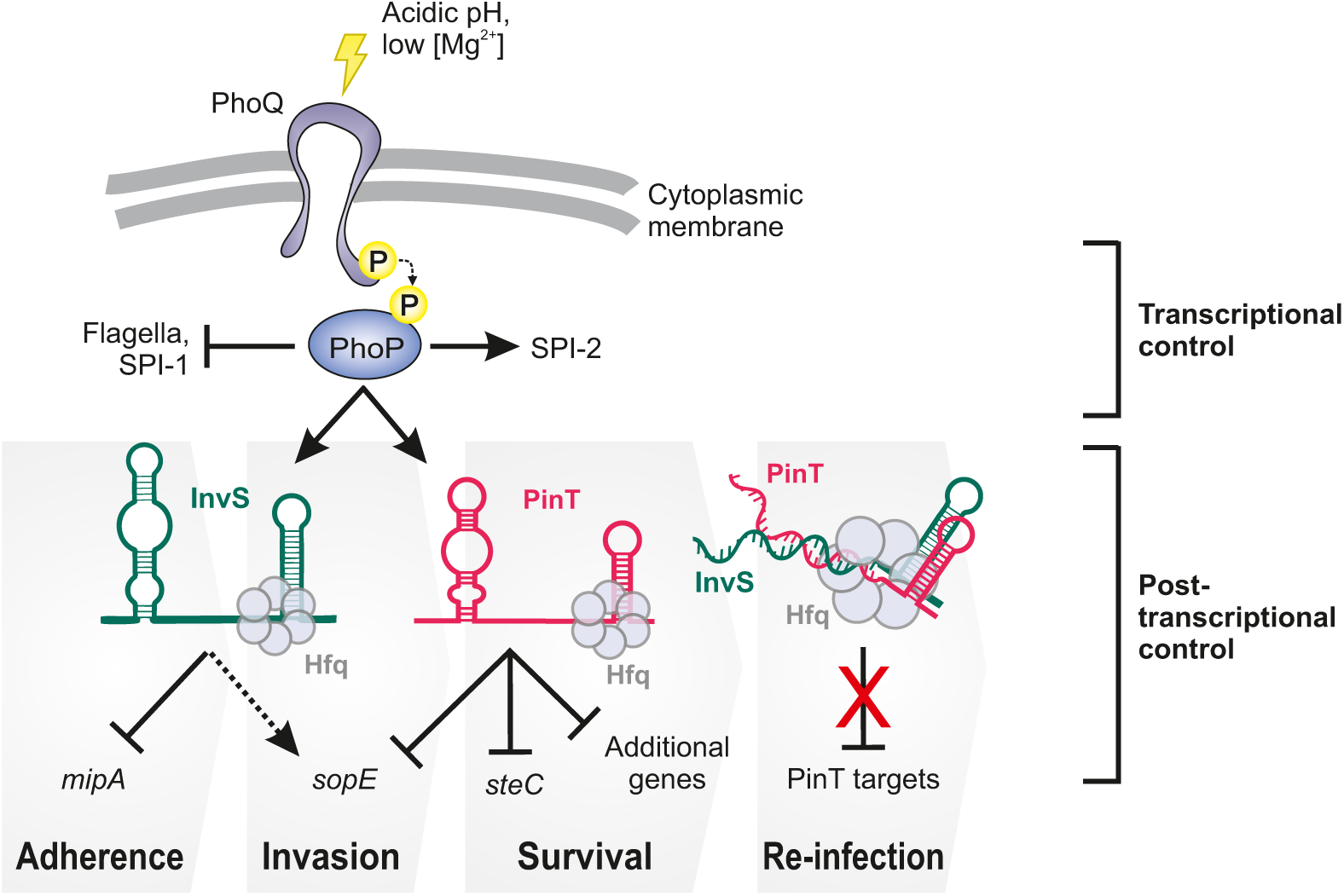
Interconnected regulatory network of two PhoP-induced sRNAs during *Salmonella* infection. The working model is based on the results presented in the current study and the existing literature (18,20,21,44).

Sponge RNAs are best studied in Enterobacteriaceae (reviewed in (36,37). Dating back 15 years ago, the first bacterial RNA sponge was discovered in *Salmonella enterica*: an intercistronic sequence within a chitosugar utilization operon that sequestered the abundant ChiX sRNA (46,63). In the following, stable processing products of a precursor tRNA (64) and a polycistronic mRNA (65) were identified as sponge RNAs in *E*. *coli* and *Salmonella*. However, the advent of techniques for global proximity ligation and sequencing of base-pairing RNAs (34,66) led to the realization that RNA sponges are far more widespread and that also established mRNA-regulating sRNAs can act as decoys for other *bona fide* sRNAs, giving rise to large post-transcriptional networks (67). Independent of their origin and mode of biogenesis, sponge RNAs generally inactivate their target sRNAs through two non-mutually exclusive (68) mechanisms: sequestering the seed region and/or promoting sRNA decay. Both mechanisms are reflected in the present example: while InvS is destabilized by PinT, both sRNAs inactivate their partner sRNA by occluding its seed region. This blurs the discrimination between regulator and target for this sRNA-sRNA interaction pair.

Taken together, this study adds another facet to the function of PinT, arguably one of the best-understood virulence-related bacterial sRNAs. Molecularly understanding the complexity of post-transcriptional virulence control circuits in bacterial pathogens bears potential to be leveraged for anti-virulence therapeutic strategies against bacterial infections (69).

## MATERIALS AND METHODS

### Bacterial strains and growth conditions

*Salmonella enterica* serovar Typhimurium strain SL1344 (JVS-1574) was used as a wild-type strain. The complete list of strains used in this study is contained in Supplementary Table 2. Bacterial cells were grown at 37°C with shaking at 220 rpm in LB medium. Where appropriate, media were supplemented with antibiotics at the following concentrations: 100 µg/ml ampicilin, 50 µg/ml kanamycin, and 20 µg/mL chloramphenicol. To investigate bacterial behavior under SPI-1 inducing conditions, cultures were initially grown in LB, as previously described, until an OD_600_ of 2.0 was achieved. For simulating SPI-2 inducing conditions, cultures were first cultivated in LB to an OD_600_ of 2.0, followed by two successive washes with Phosphate-Buffered Saline (PBS). Subsequently, the bacteria were inoculated at a ratio of 1:50 and transferred to SPI-2 medium (170 mM MES, 5 mM KCl, 7.5 mM (NH_4_)_2_SO_4_, 0.5 mM K_2_SO_4_, 1 mM KH_2_PO_4_, 8 µM MgCl_2_, 38 mM glycerol, 0.1% bacto casamino acid, pH 5.8). This cultivation was conducted at 37 °C with constant shaking at 220 rpm until the cultures reached an OD_600_ of 0.3.

### Western blot analysis

Western blots were performed following previously published procedures. In brief, bacterial cultures were collected by centrifugation for 4 min a 13,000 *g* at 4°C, and the pelleted cells were dissolved in 1x protein loading dye to a final concentration of 0.01 OD/µL. The samples were heated up for 10-15 min at 95°C, and 0.1 ODs were separated on a 12.5% polyacrylamide gel. Proteins were transferred onto a nitrocellulose membrane for 90 min at 0.34 A, using a semi-dry blotter in transfer buffer (25 mM Tris-HCl pH 8.3, 190 mM glycine, 20% methanol). Membranes were blocked with 5-10% milk for 1 h at room temperature and rinsed in 1xTBS-Tween buffer (20 mM Tris-HCl pH 7.6, 150 mM NaCl, 0.1% Tween20). After blocking, membranes were incubated with the primary antibodies (monoclonal α-FLAG, Sigma-Aldrich #F1804; polyclonal α-GroEL, Sigma-Aldrich #G6532; custom-made, rabbit, polyclonal α-MipA, Eurogentec [Supplementary Table 2]) diluted in 1x TBS-Tween buffer containing 3% BSA. The membranes were washed three times for 15 min with agitation in 1x TBS-Tween buffer. Membranes were then incubated for 1 hour at room-temperature with HRP-linked secondary antibodies (α-mouse or α-rabbit, Cell Signaling Technology #7076 or #7074 [Supplementary Table 2]), diluted in 1x TBS-Tween containing 3% BSA, and washed three times for 15 min with 1x TBS-Tween. The membranes were developed using Amersham ECL Prime reagents (GE Healthcare) and signals were detected on a LAS4000 and Imager 600 (GE Healthcare). Bands were quantified using EMBL ImageJ software.

### Salmonella infection assay

T75 cell flasks were seeded with 200,000 murine RAW264.7 macrophage cells at passage #16. On the day of the infection, macrophage cells were counted and overnight bacterial cultures were harvested to an amount that would determine a multiplicity of infection (MOI) of 50 bacteria per macrophage for all the needed flasks (# bacterial cells = [MOI x # of eukaryotic cells x # of flasks]/bacterial concentration). Before infection, harvested bacteria were opsonized with 10% of mouse serum for 20 min at room temperature. Opsonized bacteria were resuspended in RPMI media to the correct dilution for the envisaged MOI and inoculated in cell-containing flasks. Flasks were centrifuged at 250 g for 10 min at room temperature to force bacterial contact with host cells and synchronize the infection. Flasks were incubated for 30 min at 37°C to allow phagocytosis to take place. The RPMI medium from the flasks was replaced with fresh RPMI containing 100 μg/mL of gentamicin (high-gentamicin) to inhibit extracellular bacteria and flasks were incubated at 37°C for 30 min. RPMI medium was replaced once more with RPMI containing 10 μg/mL of gentamicin (low-gentamicin) and incubated for the remainder of the assay. At 20 h, flasks were washed once with ice-cold 1x PBS and incubated with 10 mL of 0.1% PBS-T to detach adhering cells. Harvested cells were either directly subjected to RIL-seq (strategy 1) or incubated for 10 min at room temperature in 0.1% PBS-T to ensure mild lysis of only the eukaryotic cells (strategy 2). In case of the latter, samples were then centrifuged at 250 g for 10 min at 4°C to separate eukaryotic cell debris from formerly intracellular bacteria and the bacteria-containing supernatant was collected and centrifuged at 4,500 g for 20 min at 4°C. The resulting bacterial pellet was snap-frozen in liquid nitrogen.

### RIL-seq

RIL-seq experiments were performed as described in (34,35) with a few modifications to adapt the protocol to *Salmonella*. Briefly, *Salmonella* strains carrying a wild-type or a 3x FLAG-tagged copy of *hfq* were grown in LB medium to an OD_600_ of 2.0. The samples were cross-linked under a 256 nm UV light source and pelleted in ice-cold 1x PBS. Pellets were lysed in NP-T buffer (50 mM NaH_2_PO_4_, 300 mM NaCl, 0.05% Tween, pH 8.0) supplemented with protease inhibitor (1:200) and RNase inhibitor (final concentration of 0.1 U/µL). Lysates were incubated with anti-FLAG antibody-bound protein A/G magnetic beads for 2 h at 4°C under continuous rotation, followed by three washing steps with lysis buffer. Beads were treated with an RNase A/T1 mix for 5 min at 22°C in an RNase inhibitor-free lysis buffer. Samples were washed three times with lysis buffer supplemented with 3.25 µL of SuperaseIN. The trimmed ends of RNAs were cured by PNK treatment for 2 h at 22°C with agitation, followed by two washing steps at 4°C. Proximal Hfq-bound RNAs were ligated with T4 RNA ligase I in the following reaction: 8 µL T4 ligase buffer, 7.2 µL DMSO, 0.8 µL ATP (100 mM), 32 µL PEG 8000, 1.2 µL RNase inhibitor, 23.6 µL of water, 140 U of T4 RNA ligase I. Samples were incubated overnight at 22°C with agitation, followed by three steps of washing with lysis buffer at 4°C. The RNAs were eluted from beads with a proteinase K digestion for 2 h at 55°C followed by LS Trizol extraction, as per manufacturer instructions. Purified RNA was resuspended in 7 µL of nuclease-free water and quality-controlled on a Bioanalyzer Pico RNA chip prior to cDNA library preparation.

Library preparation was conducted using the sRNA NEBNext kit for Illumina, with few modifications. Briefly, 3 µL of RNA were mixed with 1 µL of 3’ SR Adaptor (1:10 diluted) and incubated at 70°C for 2 min. 6.5 µL of 3’ ligation mix (5 µL of 3’ ligation buffer, 1.5 µL of 3’ enzyme mix) were added to each tube and incubated at 25°C for 1 h. 2.75 µL of SR mix (2.5 µL of water, 0.25 µL of SR RT primer) were added. The reactions were incubated in three sequential steps (75°C for 5 min, 37°C for 15 min, 25°C for 15 min). Pre-denatured 5’ adaptor (1:10 diluted) was added to each tube together with 5’ ligation mix (0.5 µL of 5’ ligation reaction buffer, 1.25 µL 5’ ligation enzyme mix) and samples were incubated at 25°C for 1 hour. First-strand cDNA synthesis was initiated by adding 5 µL of cDNA mix (4 µL first-strand buffer, 0.5 µL murine RNase inhibitor, 0.5 µL SuperScript II RT). The reactions were incubated at 50°C for 1 h followed by 15 min at 70°C. 10 µL of cDNA were PCR-amplified with barcoded NEB index primer and SR primer in a 50 µL reaction (25 µL LongAmp Taq 2x mix, 12.5 µL nuclease-free water, 1.25 µL SR primer, 1.25 µL index primer). The PCR cycling program was set as follows: 30 sec at 94°C initial denaturation, 15 sec at 94°C, 30 sec at 62°C, 70°C for 18-20 cycles, and a final elongation for 5 min at 70°C. PCR products were purified using AMPure XL beads and checked on a DNA Bioanalyzer to estimate the size distribution and amount of DNA fragments. Amplified cDNAs from different samples were pooled in equimolar ratio and sequenced in paired-end mode with 2 × 40 or 2 × 38 cycles on an Illumina NextSeq 500 platform.

Raw read pairs were quality and adapter trimmed via Cutadapt (70) v2.5 in paired-end mode using a cutoff Phred score of 20. Read pairs without any remaining bases in at least one read of a pair were discarded (parameters: --nextseq-trim=20 -m 1 -a AGATCGGAAGAGCACACGTCTGAACTCCAGTCAC -A GATCGTCGGACTGTAGAACTCTGAACGTGTAGATCTCGGTGGTCGCCGTATCATT). Processed read pairs were further analyzed using the RILseq software package (34,35) v0.82 (https://github.com/asafpr/RILseq; see https://github.com/yairra/RILseq for latest versions) installed via Bioconda (71). Here, we applied the *ad hoc* annotation of *Salmonella* Typhimurium strain SL1344 described in (15). Briefly, gene annotations from NCBI were used for the genomic features such as tRNAs, rRNAs and CDSs. Transcriptional units were defined based on the TSS annotation of (2) and Rho-independent terminator predictions using RNIE (72). The BioCyc annotation (73) was used for the plasmids. In the first step, read pairs were aligned to the *Salmonella enterica* subsp. *enterica* serovar Typhimurium str. SL1344 reference genome (RefSeq assembly accession: GCF_000210855.2) using the map_single_fragments.py script with default parameters. BWA (74)v0.7.17 was used for read alignment with the RILseq software. A transcript file (TF) with transcriptional unit annotations was generated and included for chimera mapping based on the same reference genome as above by applying the map_chimeric_fragments.py script separately to each of the previously generated BAM files. For this, default parameters were used except for setting the parameter -t TF to incorporate the transcript file in the analysis. For subsequent analyses, chimeras from conditions with two replicates were unified as described in (35). Finally, overrepresented interacting regions were identified using the RILseq_significant_regions.py script with default parameters except for including a BioCyc data folder for *Salmonella* (--bc_dir), the chromosome ID mapping --BC_chrlist NC_016810.1,NC_016810,NC_017718.1,NC_017718,NC_017719.1,NC_017719,NC_017720.1,NC_0 17720) and parameters for excluding rRNA interactions (--ribozero --rrna_list rRNA,RRNA). The resulting significant interacting regions (S-chimeras) were reannotated using the *Salmonella ad hoc* annotation described above. Summary statistics for sequencing and mapping were calculated as suggested in (35) and can be found in Supplementary Table 1. A filter of ≥40 was applied to the number of chimeric fragments supporting the S-chimeras. From the resulting, filtered dataset, RIL-seq-related figures were generated as previously described (30). BED files for visualization of the InvS-*mipA* interaction based on aligned chimeric fragments in the IGB genome browser (Fig. 3B) were generated using the generate_BED_file_of_endpoints.py script from the RILseq software.

### Translational GFP reporter assays

Strains carrying a superfolder-GFP reporter plasmid (51) were grown as described above and streaked on LB-agar followed by UV-exposure to visualize GFP expression. For the fluorescence reporter assay, specific strains were grown overnight in LB or SPI-2 media and the next day, subcultures with a starting OD_600_ of 0.02 were transferred to a 96-well plate (BRAND, # 781971). GFP signal intensities and cell densities were measured every 10 min for 24 hours, while the plate was continuously shaking (200 rpm) at 37°C in a BioTek Synergy H1 Plate Reader. Wells containing pure media were included and their background fluorescence and opacity used as blanks.

### In-vitro transcription and RNA labelling

200 ng of a DNA fragment PCR-amplified from *Salmonella* genomic DNA was used as a template in a T7 transcription reaction using the MEGAscript T7 Transcription kit (Thermo Fisher Scientific). The size and integrity of RNA was confirmed on a denaturing polyacrylamide gel. RNA bands were excised from the gel and eluted in RNA elution buffer (0.1 M sodium acetate, 0.1 % SDS, 10 mM EDTA at 4°C overnight), isolated with Phenol:Clorophorm:Isoamyl (P:C:I) and precipitated in EtOH. 50 pmol of RNA was dephosphorylated with 10 units of calf intestinal phosphatase (CIP, New England Biolabs) in a 50 μL reaction at 37°C for 1 h. CIP-treated RNA was extracted with P:C:I and EtOH precipitated. 20 pmol of the dephosphorylated RNA was 5’-labelled with 2 μL of ^32^P-γ-ATP (10 μCi/μL) using 1 unit of T4 polynucleotide kinase (Thermo Fisher Scientific) for 1 h at 37°C in a 20 μL reaction. RNA was purified from unincorporated nucleotides with microspin G-50 columns (GE Healthcare) according to the manufacturer’s instructions.

### Electrophoretic mobility shift assay

0.04 pmol of radio-labelled RNA was used for each reaction mix. Labelled RNA was denatured at 95°C for 1 min and chilled on ice for 5 min. 1x structure buffer (10 mM Tris-HCl pH 7.0, 0.1 M KCl, 10 mM MgCl_2_) was added and the RNA was re-natured at 37°C for 10 min. Per reaction, 1 μg of yeast RNA was added to the mix and the labelled RNA was added to tubes containing increasing concentration of unlabelled RNA. Where applicable, 100 nM of purified Hfq were added to the reaction mix. Reactions were incubated at 37°C for 15 min, stopped by adding 5x RNA native loading buffer and separated on a native 6% polyacrylamide gels at 4°C in 0.5% TBE at constant current of 40 mA for 3-4 h. Gels were dried and signals detected on a Typhoon FLA 7000 phosphoimager and quantified with EMBL ImageJ software.

### RNA structure probing

Labeled RNAs were prepared as described for the EMSA. For the reactions, 0.4 pmol of labeled RNA were denatured as described earlier and incubated with increasing concentration of unlabeled RNA partner (InvS) for 15 min at 37°C in the presence of 1x Structure buffer and 1 µg of yeast RNA in 10 µL. The reactions were treated with 2 µL of 25 nM Lead-Acetate and incubated for 90 sec at 37°C. 12 µL of GL II RNA loading dye were added to each tube to stop the reaction. 10 µL of each sample were boiled at 95°C for 3 min, loaded on a 10% PAA 7 M Urea gel, and separated for 3 hours at 45 Watts. For the CTRL lane, 1 pmol of labeled RNA was denatured at 95°C in 10µL of water and stopped on ice with 10 µL of GL II RNA loading dye. For the OH ladder, 1 pmol of labeled RNA was denatured at 95°C for 5 min in 1x alkaline buffer in a 10 µL reaction. For the T1 ladder, 1 pmol of RNA was denatured in water for 1 min at 95°C followed by addition of RNase T1 enzyme and incubated for 3 min at 37°C. All reactions were stopped as mentioned before.

### Northern blot analysis

4 OD equivalents of Bacterial cultures were snap-frozen in liquid nitrogen after adding 0.2 vol/vol of stop solution (95% ethanol and 5% phenol). Bacterial cells were pelleted and lysed by adding 600 µL of lysozyme (0.5 mg/mL) and 60 µL of 10% SDS, followed by a 2-minute incubation at 64 °C. Subsequently, 66 µL of 3 M sodium acetate (pH 5.2) and 750 µL of phenol were added, with the mixture incubated for 6 minutes at 64 °C. After centrifugation, the aqueous phase was transferred to a PLG tube, followed by the addition of 750 µL chloroform. After a second centrifugation, the aqueous phase was collected. RNA was then precipitated overnight at −20 °C using a 30:1 mixture of ethanol and 3 M sodium acetate (pH 6.5), pelleted, washed with 75% ethanol, and resuspended in 50 µL of nuclease-free water. 5 to 10 μg of total RNA were denatured at 95°C for 5 min in RNA loading dye (95% v/v formamide, 10 mM EDTA, 0.1% w/v xylene cyanole, 0.1% w/v bromophenol blue) and separated on a gel electrophoresis with 6% polyacrylamide/7 M urea in 1x TBE buffer for 2 h at 300 V. RNA was transferred onto a Hybond-XL nylon membrane (GE Healthcare) with electro-blotting at 50 V for 1 h at 4°C. The membrane was cross-linked at 120 mJ/cm^2^ with UV light and pre-hybridized for 10 min in Rapid-Hyb buffer (Amersham). A [^32^P]-labeled oligonucleotide probe was added onto the membrane and hybridized at 42°C overnight with rotation. The membrane was washed three times for 15 min with 5x SSC/0.1% SDS (first wash), 1x SSC/0.1% SDS (second wash) and 0.5x SSC/0.1% SDS (third wash) buffers at 42°C. Air dried membranes were then exposed onto a phosphor screen and signals were visualized on a Typhoon scanner and quantified with the ImageJ software.

### RNA stability assay

In this assay, cultures in SPI-2 medium, with an optical density at OD_600_ of 0.3, were treated with rifampicin to achieve a concentration of 500 μg/mL. At specified intervals post-rifampicin treatment, samples were taken for total RNA isolation. The extracted RNA was subsequently subjected to analysis via northern blotting, utilizing the previously outlined method.

### qRT-PCR analysis

To eliminate any genomic DNA contamination, the RNA samples underwent a purification step using 0.25 U of DNase I, RNase free (Thermo Fisher Scientific, #EN0521) for every 1 µg of RNA, incubating for 45 min at 37°C. The qRT-PCR analysis was conducted utilizing the Takyon No ROX SYBR 2x Mastermix blue (Eurogentec #UF-NSMT-B0701) in accordance with the guidelines provided by the manufacturer, employing a QuantStudio 5 Real-Time PCR System (Thermo Fisher Scientific) for detection.

## ACKNOWLEDGEMENTS

We thank Sarah Reichardt for excellent technical support and the Core Unit Systems Medicine at the University of Würzburg for excellent technical support, RNA-seq data generation, and analysis. The Core Unit Systems Medicine is partly supported by the Interdisciplinary Center for Clinical Research (IZKF) at the University of Würzburg (project Z-6). This study was funded by the DFG Research Training Group GRK2157 “3D-Infect” (to J.V. and A.J.W.). Research in the Westermann lab is supported by the European Research Council (ERC Starting Grant #101040214 to A.J.W.). H.K. acknowledges funding through the HIRI graduate training program “RNA & Infection”.

## AUTHOR CONTRIBUTIONS

J.V. and A.J.W. conceived of and designed the project. H.K., G.M., and E.V. performed experiments. RIL-seq data were generated by G.M. and analyzed by T.B., G.M., and H.K. A.J.W. wrote the manuscript primarily with input/editing from H.K., G.M., and J.V. J.V. and A.J.W. secured funding and supervised the study.

## SUPPLEMENTARY ITEM LEGENDS

**Supplementary Figure S1.**
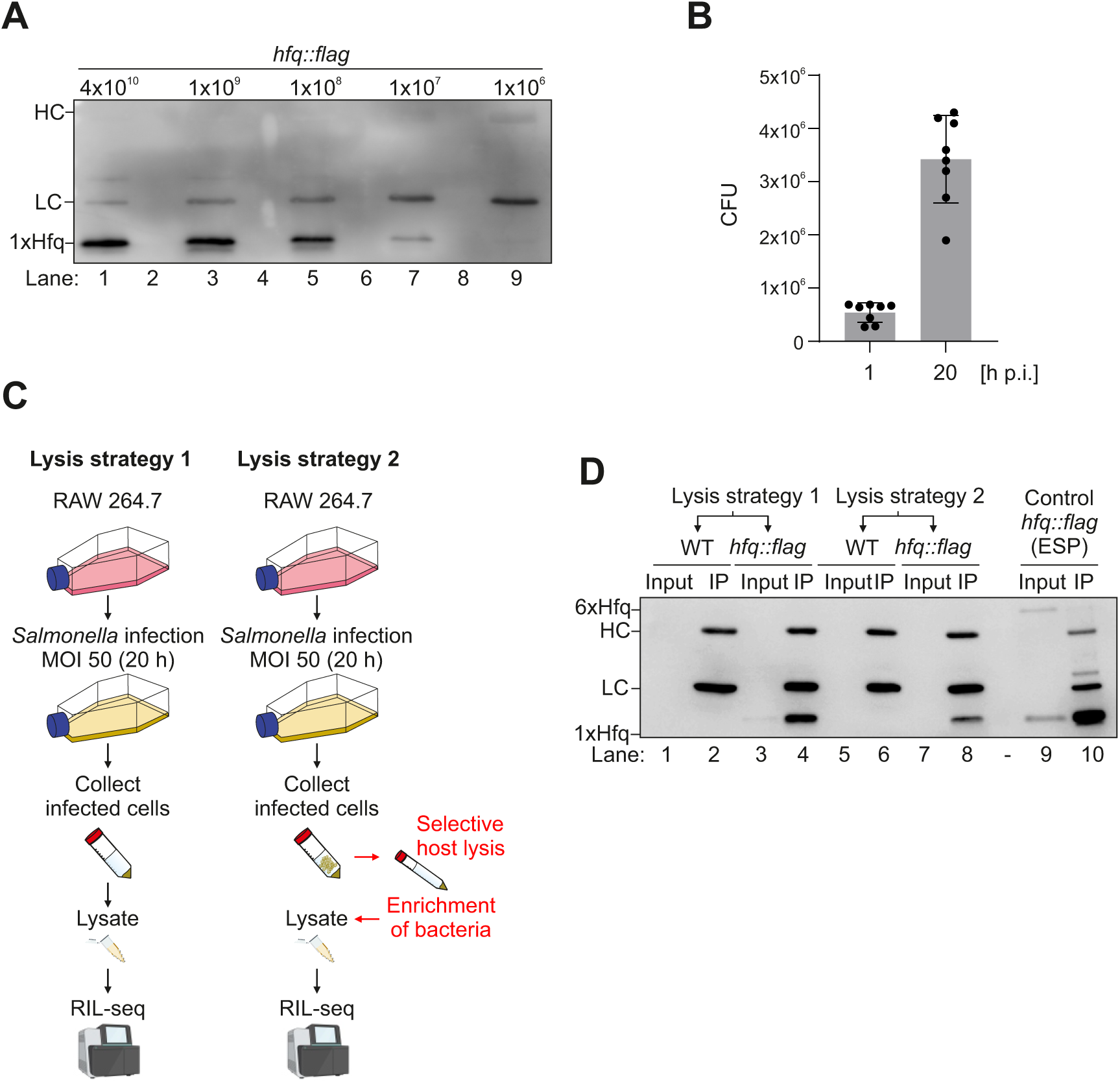
Optimization of the intramacrophage RIL-seq approach. **A,** Western blot showing the minimum number of bacteria necessary to successfully pulldown the FLAG-tagged Hfq protein. **B,** RAW264.7 macrophages were infected at a multiplicity of infection (MOI) of 50 with wild-type *Salmonella* and the intracellular bacteria at 1 and 20 hours post infection (p.i.) were enumerated by plating assays. **C,** Two lysis strategies were evaluated for intramacrophage RIL-seq. For both of them, infected macrophages were harvested at 20 h p.i. and pelleted by centrifugation. These pellets were either directly subjected to lysis (strategy 1) or enriched for intracellular bacteria (strategy 2). In case of the latter, harvested macrophages were incubated in 0.1% PBS-Triton X-100 for 10 min at room temperature for selective lysis of only eukaryotic cells. The resulting lysate was centrifuged at 250 g for 10 min, 4°C, to pellet host cells remnants, while the released bacteria were retained in the supernatant. The supernatant was collected and centrifuged at 4,500 g for 20 min, 4°C, to also pellet the bacteria, which were then snap-frozen. **D**, Although more Hfq protein could be immunoprecipitated with the “lysis1” method, a substantially higher amount of contaminant eukaryotic RNA was present in the sample. Therefore, the percentage of reads that successfully mapped to the *Salmonella* genome was substantially lower in “lysis1” compared to “lysis2”, 10% and 34%, respectively.

**Supplementary Figure S2.**
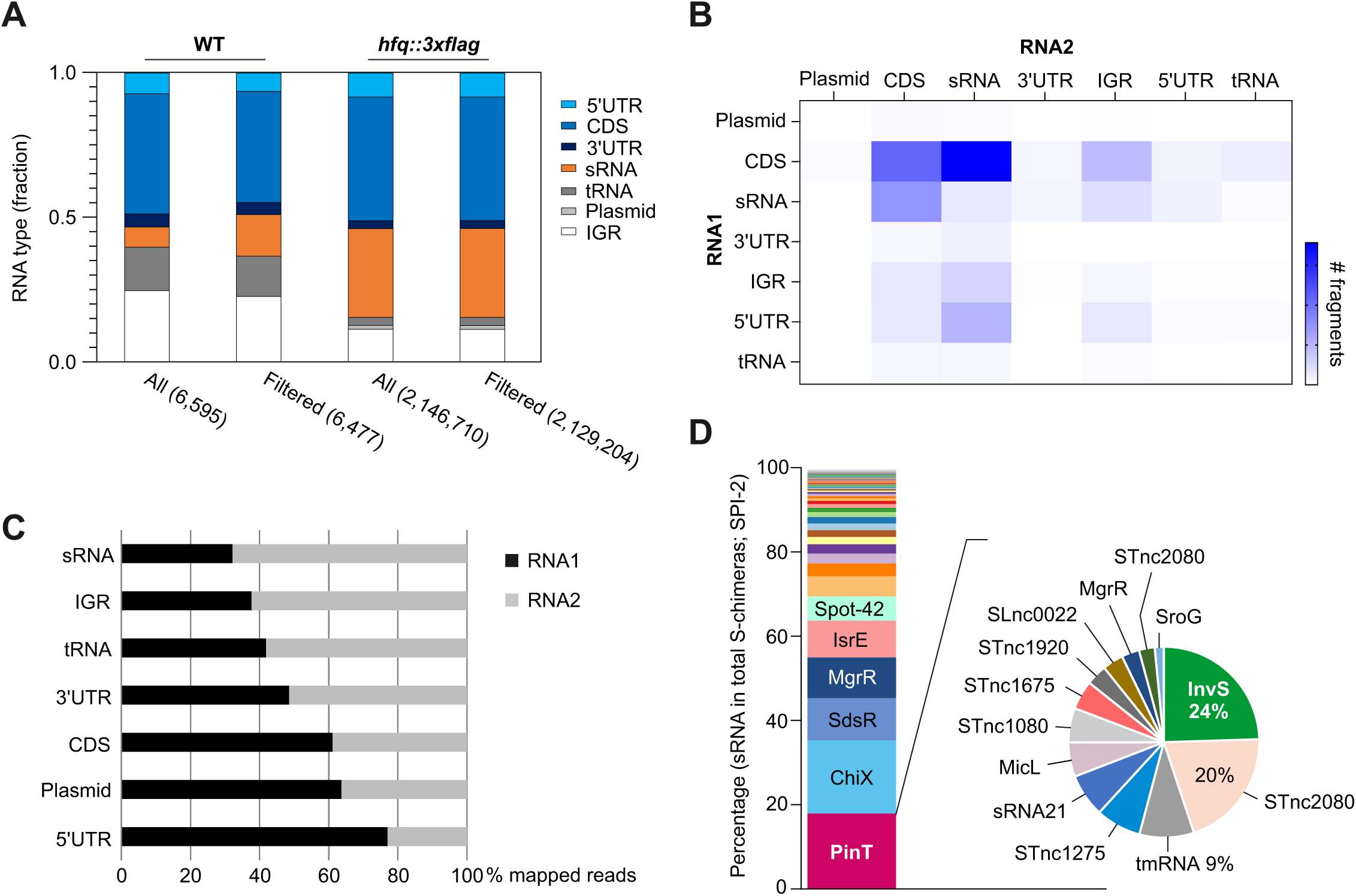
Global results of the SPI-2 RIL-seq experiment. **A,** Relative frequency of each RNA type for significant chimeric fragments (number of fragments in S-chimeras; unfiltered [all] or filtered for ≥40 interactions) within the RIL-seq dataset derived from the wild-type and *hfq::3×FLAG* strains in the SPI-2 condition. UTR, untranslated region; CDS, coding sequence; sRNA, small RNA; tRNA, transfer RNA; plasmid, plasmid-encoded transcript; IGR, intergenic region. **B,** Heat map of the number of chimeric fragments corresponding to abundance-filtered S-chimeras according to the position of the RNA within the chimera (RNA1 = first read, RNA2 = second read). **C,** Chimeric fragments in RNA1 vs. RNA2 for each RNA type in the SPI-2 RIL-seq dataset. **D,** Distribution of all sRNAs in S-chimeras of SPI-2-inducing condition and PinT-sRNA interactome based on RIL-seq data. Related to Fig. 1.

**Supplementary Figure S3.**
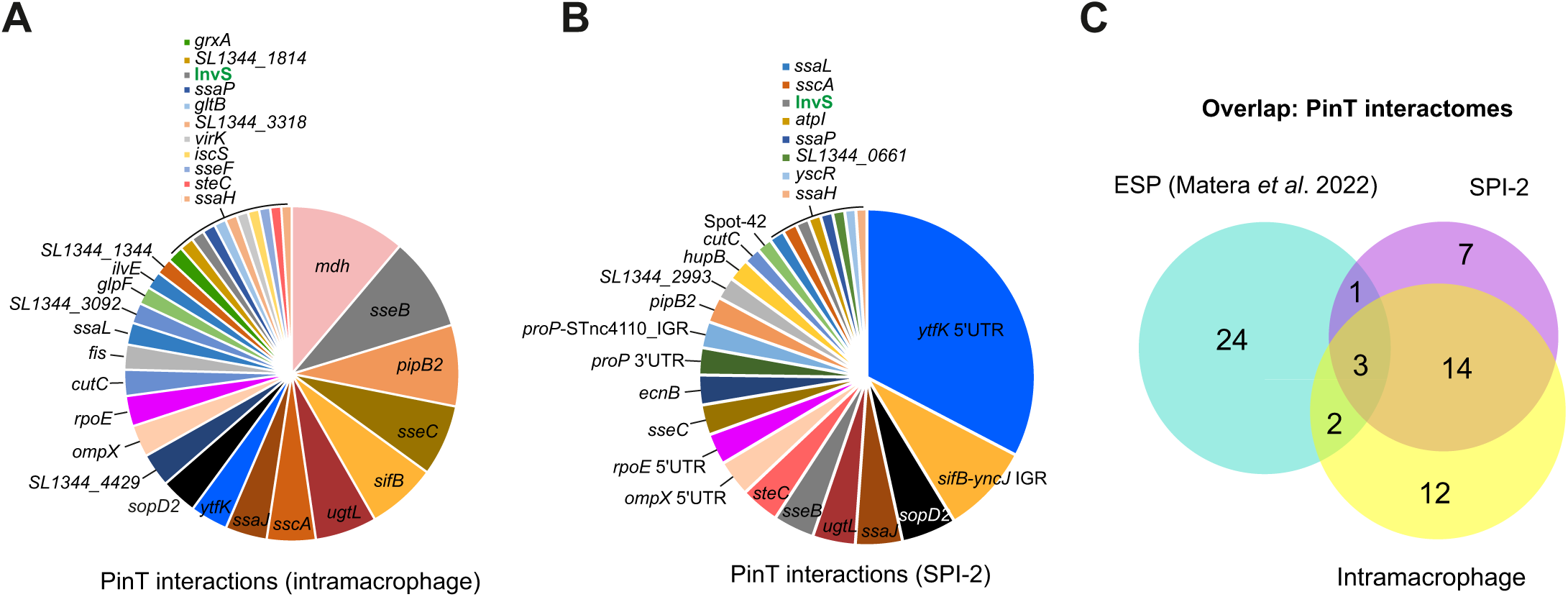
PinT targets recovered by RIL-seq under diverse conditions. **A,** PinT interactome inside macrophages. **B,** PinT interactome under the SPI-2 inducing condition. **C,** Venn diagram illustrating the overlap of the PinT interactomes derived from three distinct *Salmonella* Hfq RIL-seq data sets: intramacrophage (this study), *in vitro*-growth in SPI-2-inducing medium (this study), and growth in rich medium to early stationary phase (ESP) (30). The diagram displays the number of unique and overlapping interactions identified, highlighting shared and condition-specific interactors of PinT. Related to Fig. 2G, H.

**Supplementary Figure S4.**
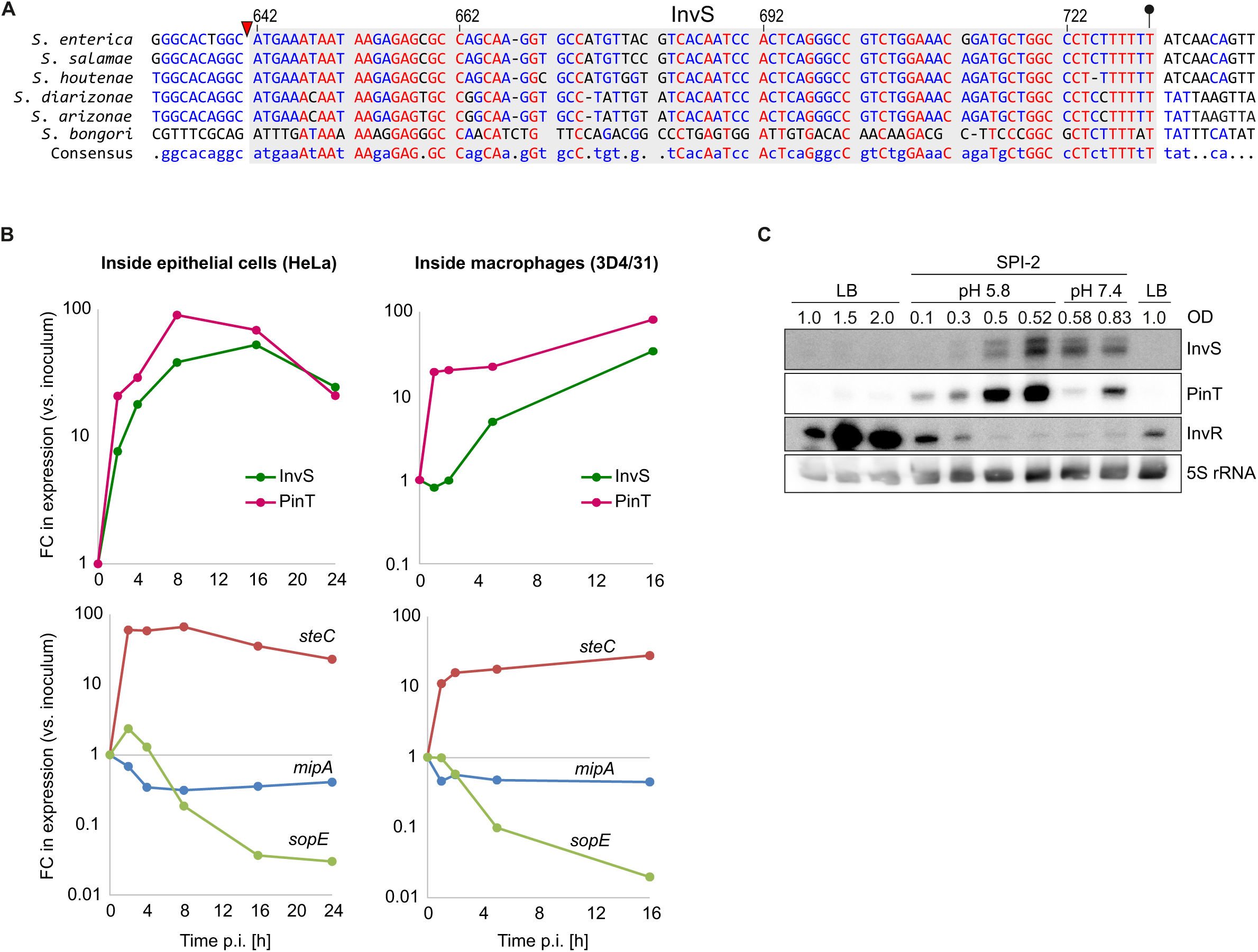
Expression kinetics of InvS and PinT and of their target mRNAs by intracellular *Salmonella*. **A,** Genomic locus of *SL0083A-srfN-invS*. **B,** sRNA expression (upper) and their target mRNA expression (lower) during the infection of epithelial cells (left) or of macrophages (right) relative to the corresponding expression levels in the bacterial inoculum. The original RNA-seq data stem from (18). **C,** In-vitro SPI-1/SPI-2 transition assay. For each transition between LB and SPI-2-inducing medium the bacteria were inoculated at a ratio of 1:50, to ensure consistency and synchronized growth.

**Supplementary Figure S5:**
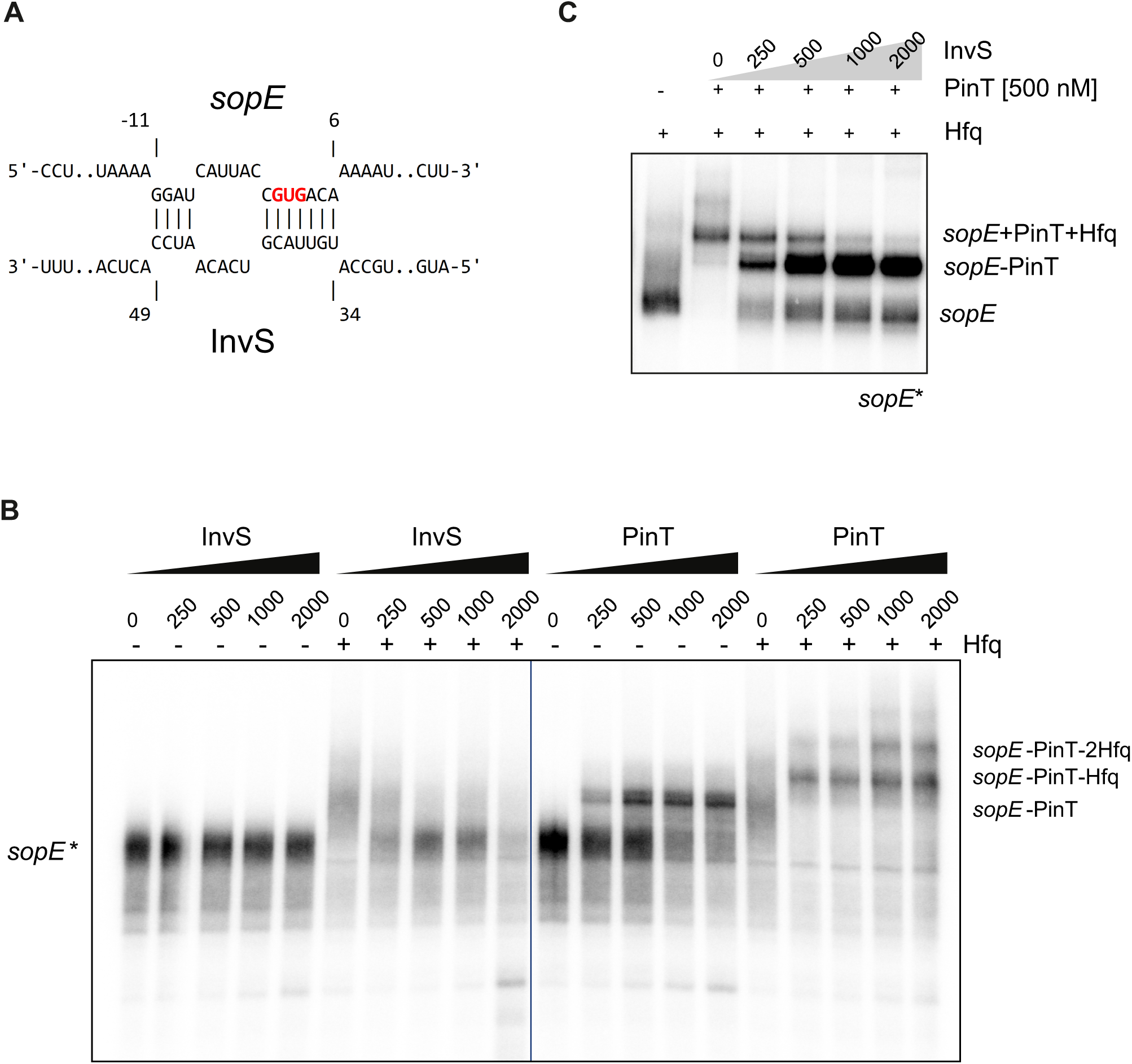
Evaluation of *sopE* mRNA as an InvS target candidate. **A,** IntaRNA-derived prediction of InvS base-pairing to the translation initiation region of *sopE* mRNA. Numbers refer to the position relative to the 5’ end in case of InvS and relative to the start codon of *sopE*, which is highlighted in red. **B,** The radioactively labeled 5′ fragment of *sopE* does not bind to InvS but binds to PinT both independently and in the presence of the RNA chaperone Hfq, forming a stable complex. **C,** InvS competes with PinT-*sopE* duplexes for Hfq binding. Three-component EMSA was performed by increasing concentrations of InvS and continuously present Hfq (100 nM).

**Supplementary Figure S6:**
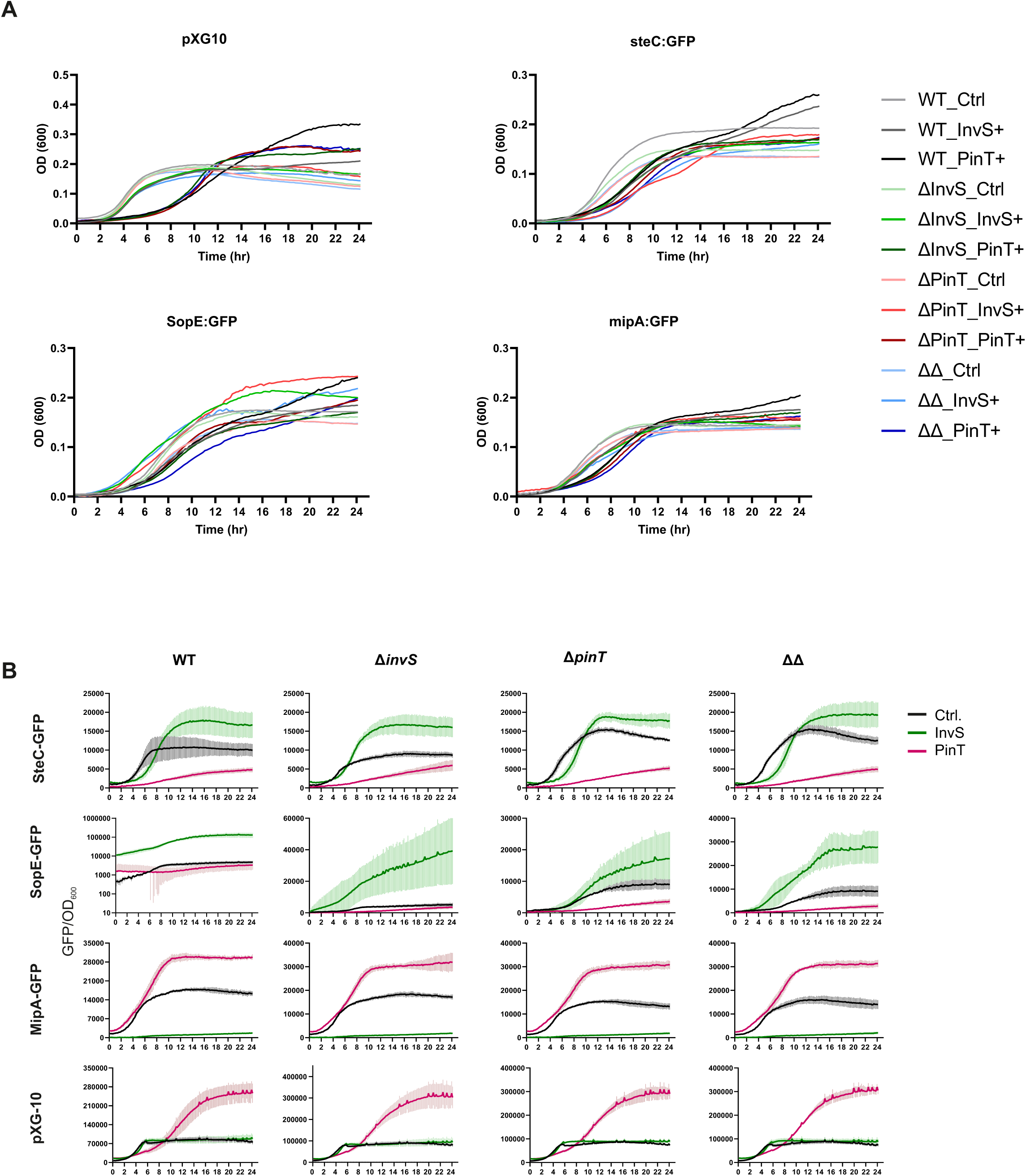
Dual plasmid reporter assay of InvS and PinT targets in defined mutant backgrounds and corresponding growth kinetics. **A,** To monitor bacterial growth and sRNA target expression, cultures were diluted in SPI-2 MM and dispensed into a 96-well plate. Optical density was measured at 10 minute intervals throughout a 24 hour incubation. Data represent the mean of three biological replicates, each with two technical replicates. Standard deviations (SD) were omitted for clarity in visualizing the mean growth curves. **B,** A dual-plasmid reporter assay was conducted to evaluate the regulation of InvS and PinT targets in defined mutant backgrounds. Each strain was transformed with either a control plasmid or plasmids constitutively expressing InvS or PinT. Optical density at 600 nm (OD₆₀₀) and sfGFP fluorescence were measured at regular intervals over a 24-hour period using a plate reader. GFP expression was normalized by calculating the GFP/OD₆₀₀ ratio for each well. Data represent the mean of three independent biological replicates, each measured in duplicate.

**Supplementary Table 1: RIL-seq data.** Sheet 1: counts of sequenced and mapped fragments in individual datasets, as identified through RIL-seq. Sheet 2: significant interactions unified over the two intramacrophage RIL-seq experiments (following lysis strategy #2). Sheet 3: significant interactions unified over the two SPI-2 RIL-seq libraries.

**Supplementary Table 2: Bacterial strains, plasmids, oligonucleotides, and antibodies used in this study.**

